# Initial insights into the genetic epidemiology of SARS-CoV-2 isolates from Kerala suggest local spread from limited introductions

**DOI:** 10.1101/2020.09.09.289892

**Authors:** Chandni Radhakrishnan, Mohit Kumar Divakar, Abhinav Jain, Prasanth Viswanathan, Rahul C. Bhoyar, Bani Jolly, Mohamed Imran, Disha Sharma, Mercy Rophina, Gyan Ranjan, Beena Philomina Jose, Rajendran Vadukkoot Raman, Thulaseedharan Nallaveettil Kesavan, Kalpana George, Sheela Mathew, Jayesh Kumar Poovullathil, Sajeeth Kumar Keeriyatt Govindan, Priyanka Raveendranadhan Nair, Shameer Vadekkandiyil, Vineeth Gladson, Midhun Mohan, Fairoz Cheriyalingal Parambath, Mohit Mangla, Afra Shamnath, Indian CoV2 Genomics & Genetic Epidemiology (IndiCovGEN) Consortium, Sridhar Sivasubbu, Vinod Scaria

## Abstract

Coronavirus disease 2019 (COVID-19) rapidly spread from a city in China to almost every country in the world, affecting millions of individuals. Genomic approaches have been extensively used to understand the evolution and epidemiology of SARS-CoV-2 across the world. Kerala is a unique state in India well connected with the rest of the world through a large number of expatriates, trade, and tourism. The first case of COVID-19 in India was reported in Kerala in January 2020, during the initial days of the pandemic. The rapid increase in the COVID-19 cases in the state of Kerala has necessitated the understanding of the genetic epidemiology of circulating virus, evolution, and mutations in SARS-CoV-2. We sequenced a total of 200 samples from patients at a tertiary hospital in Kerala using COVIDSeq protocol at a mean coverage of 7,755X. The analysis identified 166 unique high-quality variants encompassing 4 novel variants and 89 new variants identified for the first time in SARS-CoV-2 samples isolated from India. Phylogenetic and haplotype analysis revealed that the circulating population of the virus was dominated (94.6% of genomes) by three distinct introductions followed by local spread, apart from identifying polytomies suggesting recent outbreaks. The genomes formed a monophyletic distribution exclusively mapping to the A2a clade. Further analysis of the functional variants revealed two variants in the *S* gene of the virus reportedly associated with increased infectivity and 5 variants that mapped to five primer/probe binding sites that could potentially compromise the efficacy of RT-PCR detection. To the best of our knowledge, this is the first and most comprehensive report of genetic epidemiology and evolution of SARS-CoV-2 isolates from Kerala.

## INTRODUCTION

The COVID-19 pandemic has seen a widespread application of genomic approaches to understand the epidemiology and evolution of SARS-CoV-2. The accelerated efforts to sequence genomes of clinical isolates of SARS-CoV-2 from across the world picked up pace following the initial genome sequencing of the virus from a patient in Wuhan, the epicenter for the pandemic [1]). As the virus evolves through the accumulation of mutations, it has split into major lineages with strong geographical affinities [2]. The availability of the genome sequences in the public domain has provided a unique view of the introduction, evolution, and dynamics of SARS-CoV-2 in different parts of the world [3].

A number of approaches have emerged for rapid and scalable sequencing of SARS-CoV-2 from clinical isolates. This includes direct shotgun approaches as well as targeted amplicon-based and targeted capture-based approaches [4–6]. Sequencing based approaches provide a unique opportunity for high fidelity of detection and for understanding the genetic epidemiology of SARS-CoV-2 [7]. Additionally, the genetic variants could offer insights into the mutational spectrum, evolution, infectivity, and attenuation of the virus [8,9]. Additional analyses on genomic variants have also provided useful insights into the efficacy of primer/probe-based diagnostic assays as well as immune epitopes and resistance to antisera [10,11]. An approach for high-throughput multiplex amplicon sequencing of SARS-CoV-2 has been previously reported from our group [7].

Kerala is a unique state in India with a population of 35 million people and extensively connected with the global populations through over 1.6 million expatriates, apart from being a traditional trade post and a global tourist destination. The state is therefore in a distinct position, affected by local as well as global epidemics. In fact, the first identified case of COVID-19 in India was from Kerala, early in the epidemic. The patient had traveled from Wuhan, China [12,13]. The initial genomic identity of the virus was also established which mapped to the B superclade of SARS-CoV-2 [14]. Further introductions into the state during the later days of the pandemic through international and regional travel could have contributed to the spread of the epidemic in the state and the pool of circulating genetic lineages or clades. While a number of studies on the genetic epidemiology of SARS-CoV-2 from different states in India have emerged [14–17], there has been a paucity of information on the genetic architecture and epidemiology of SARS-CoV-2 isolates in the state of Kerala.

We intended to fulfill the gap in knowledge on the identity of the circulating genetic lineages/clades contributing to the epidemic in the state of Kerala. To this end, we employed a high-throughput sequencing-based approach for the genetic epidemiology of SARS-CoV-2. To the best of our knowledge, this is the first comprehensive overview of the genetic architecture of SARS-CoV-2 isolates from the state of Kerala.

## MATERIALS AND METHODS

### Samples and RNA isolation

The institutional Human Ethics Committee approved the project (GMC KKD/RP2020/IEC438). RNA samples were isolated from nasopharyngeal/oropharyngeal swabs of patients presenting to Government Medical College, Kozhikode, a major tertiary care center in Kerala. RNA extraction was done using MagMax Viral/Pathogen Nucleic Acid Isolation kit in Thermo Scientific KingFisher Flex automated extraction system according to the manufacturer’s instructions. All the RNA samples were transferred within 72 hours of collection at a cold temperature (2-8°C) and were stored at −80°C until further processing.

### Sequencing and Data Processing

Sequencing was performed using the COVIDSeq protocol as reported previously [7]. Briefly, this protocol involved multiplex amplicon sequencing on the Illumina NovaSeq platform. The base calls generated in the binary base call (BCL) format were demultiplexed to FASTQ reads using bcl2fastq (v2.20). For reference-based assembly, we followed a previously defined protocol from Poojary et al. [18]. As per the protocol, the quality control of FASTQ reads was performed using Trimmomatic (v0.39) at a Phred score of Q30 [19] with adapter trimming. These reads were further aligned to the severe acute respiratory syndrome 2 (SARS-CoV-2) Wuhan-Hu-1 reference genome (NC_045512.2) using HISAT2-2.1 [20]. The human reads were removed using SaMtools (v1.10) [21]. The samples with coverage >99% and <5% unassigned nucleotides underwent variant calling and consensus sequences generation using VarScan (v2.4.4) [22] and SaMtools (v1.10) [21], bcftools (v1.10.2), and seqtk (v 1.3-r114) [23] respectively.

### Variant Annotation and Comparison with Existing Datasets

Variants were annotated using ANNOVAR [24] employing a range of custom annotation datasets and tables. All the variants identified were systematically compared with a compendium of other Indian and global variants. A total of 93,995 complete SARS-CoV-2 genomes deposited in the Global Initiative on Sharing All Influenza Data (GISAID) database till September 1, 2020 were used for comparative analysis. Summary of the sample details along with their originating and submitting laboratories are provided in **Supplementary Table 1**. Viral genomes with a pairwise alignment ≥ 99% and gaps < 1% with the reference genome (NC_045512.2) were considered for further variant calling using SNP-sites [25]. Genetic variants compiled from a total of 1,855 high-quality genomes from India and 32,286 global genomes were considered for analysis.

### Phylogenetic Analysis

Phylogenetic analysis was performed according to the pipeline provided by Nextstrain [26]. The dataset of 2,476 complete SARS-CoV-2 genomes deposited in the GISAID database from India was used for the analysis **Supplementary Table 2**, along with 113 genomes from the current study which have 99% coverage and at least 98% pairwise alignment with the reference genome (NC_045512.2). Genomes having more than 5% Ns or missing dates of sample collection were excluded from the analysis. The phylogenetic tree was constructed and refined to a molecular clock phylogeny using the Augur framework provided by Nextstrain and was visualized using Auspice. The Phylogenetic Assignment of Named Global Outbreak LINeages (PANGOLIN) package was used to assign lineages to the genomes from this study [27]. The lineages were visualized and annotated on the phylogenetic tree using iToL [28].

### Haplotype Analysis

For haplotype analysis, the genomes were aligned to the Wuhan-Hu-1 (NC_045512.2) reference genome using MAFFT [29] and problematic genomic loci (low coverage, high sequencing error rate, hypermutable and homoplasic sites) were masked from the alignment [30]. The aligned sequences were imported into the DNA Sequence Polymorphism tool (DnaSP v6.12.03) [31] to generate haplotypes. A TCS haplotype network [32] for the genomes was constructed using the Population Analysis with Reticulate Trees software (POPART v 1.7) [33]. Times to the most recent common ancestor (tMRCA) for the haplogroups were computed following the Bayesian Markov chain Monte Carlo (MCMC) method using BEAST v1.10.4 [34]. The analysis was performed using a coalescent growth rate model along with a strict molecular clock and the HKY+Γ substitution model with gamma-distributed rate variation (gamma categories=4). MCMC was run for 50 million steps. The output was analyzed in Tracer v1.7.1 [35] and burn-in was adjusted to attain an appropriate effective sample size (ESS).

### Functional SARS-CoV-2 variants

Further, we have evaluated the SARS-CoV-2 variants based on their functional relevance. We curated a comprehensive compendium of SARS-CoV-2 variants of functional relevance as well as variants that are associated with increased infectivity and attenuation of SARS-CoV-2 from literature and preprint servers. The variants were systematically annotated and mapped to the reference genome coordinates and their respective amino acid changes. This variant compendium encompassed about 337 variants curated from 35 publications. The variants in this study were compared with the genomic variants generated using bespoke scripts.

### Variant effect on RT-PCR efficacy

We were also interested to evaluate the effect of SARS-CoV-2 variants on the efficacy of RT-PCR detection. We took a compiled list of 132 primer/probe sequences widely used in the molecular detection of SARS-CoV-2 around the globe [11]. In our analysis, we mapped the SARS-CoV-2 genetic variants obtained from Kerala genomes to the 132 primer or probes sequence and calculated the melting temperature (Tm) of the mutant with the wild type sequence. The length of primers in the curated list is greater than 13 nucleotides. The formula applied for calculating melting temperature is Tm=64.9 +41*(yG+zC-16.4)/(wA+xT+yG+zC) where w, x, y, and z are the number of A, T, G, and C nucleotides respectively [36]. **Figure 1** summarises the schematic for the overall data analysis.

**Figure 1.**
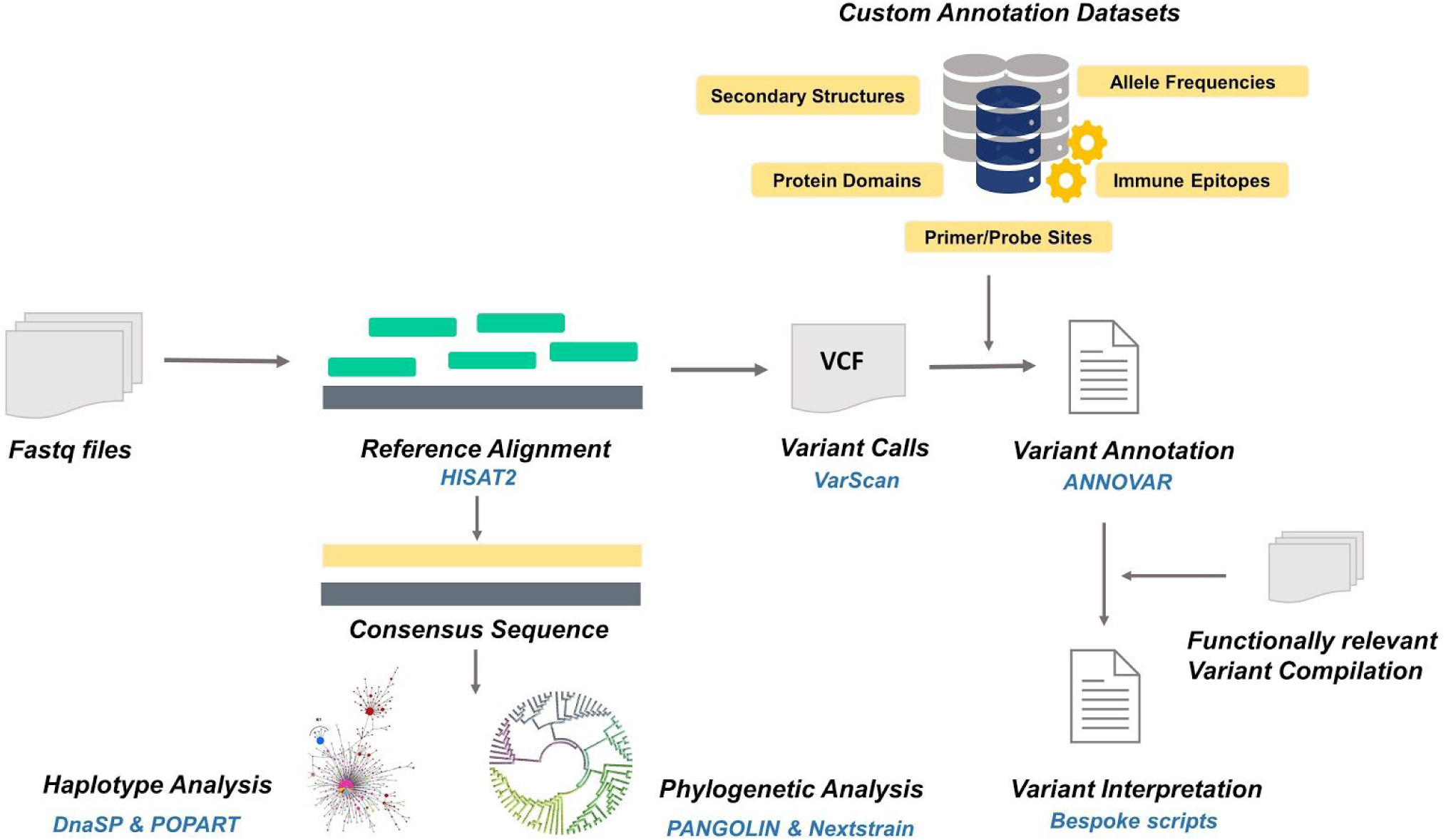
Schematic overview of the data analysis workflow comprising sequence alignment, variant calling, variant annotation, phylogenetic, and haplotype analysis employed in the study.

## RESULTS

### Sequencing and Data Processing

A total of 200 isolates of SARS-CoV-2 from Kerala were processed for genome sequencing. The genomes were sequenced using COVIDSeq protocol [7] and generated approximately 8.1 million raw reads per sample. The reads were subjected to quality control and resulted in approximately 7.5 million reads per sample, of which around 6.4 million reads per sample aligned to the SARS-CoV-2 reference genome (NC_045512.2). The reads had a mapping percentage of 84.93% and mean 7,755X coverage. The data has been summarized in **Supplementary Table 3** and the mean coverage of the sample across the amplicons has been represented in **Figure 2**.

**Figure 2:**
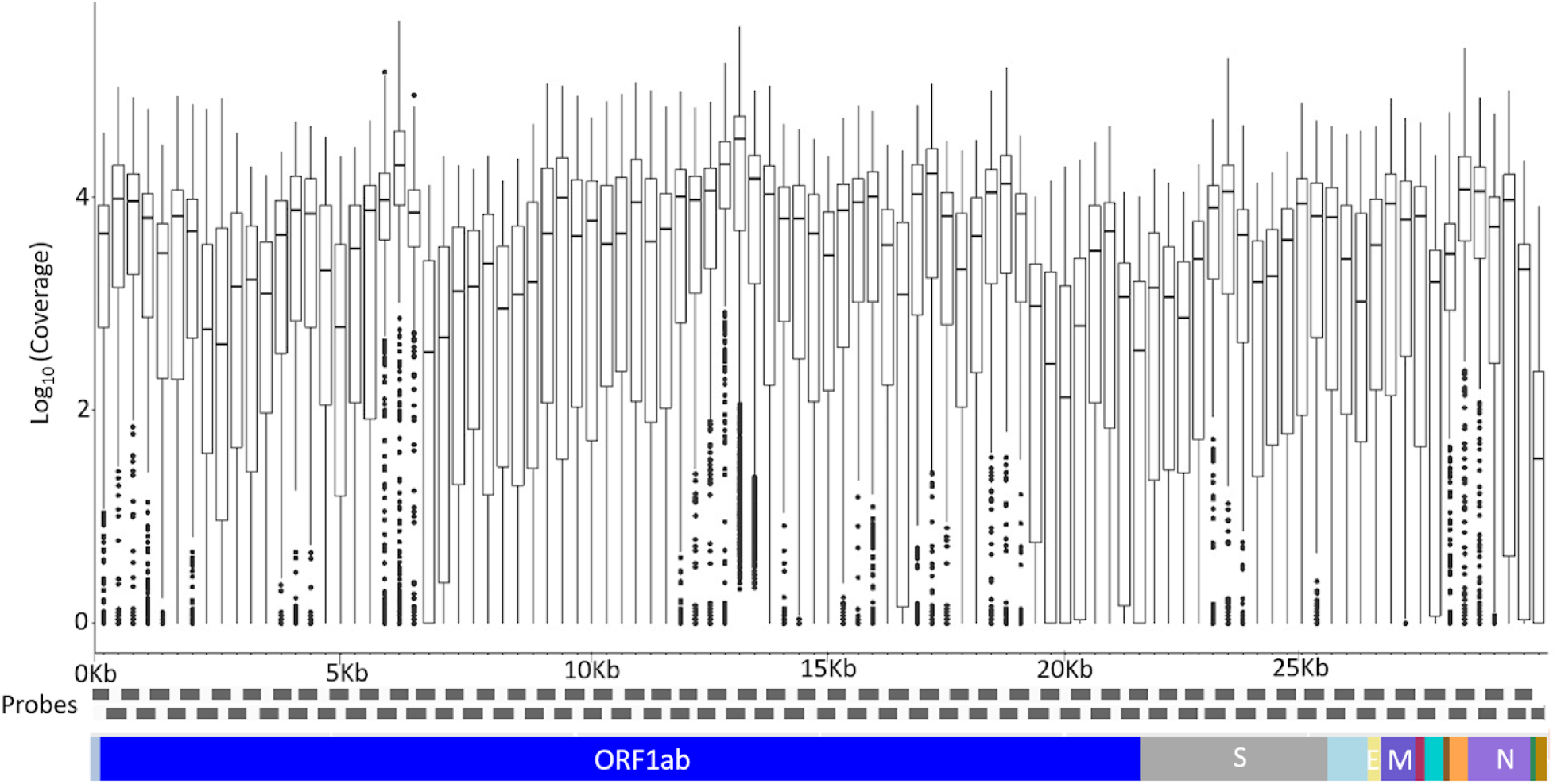
The mean coverage of the SARS-CoV-2 genomes across the amplicons of the COVIDSeq

### Variant Annotation and Comparison with Existing Datasets

Of the 200 isolates of SARS-CoV-2 sequenced, a total of 179 samples had > 99% coverage and <5% unassigned nucleotides across the genome. These samples were further processed for variant calling and consensus generation. Our analysis identified a total of 195 unique variants, with a median variant count of 12 per sample. Variant quality has been ensured with the average variation percentage across genomes ≥ 50. Of the total 195 unique variants, 166 were categorized as high-quality variants. The detailed information on variant quality is provided in **Supplementary Table 4**. The distribution of variants across the SARS-CoV-2 genomes used in the study was analyzed. Also, the proportional distribution of variants for every 100 bps across the genome was calculated and compared among various datasets. Variant distribution across genomes and comparison of variant proportions across genome datasets are represented in **Figure 3**. Out of the 166 high-quality unique variants, 4 variants were found to be reported for the first time in the global compilation of variants **Table 1**. We have also added 89 new variants (2.61%) to the Indian repertoire of genetic variants. Details of these variants are systematically compiled in **Supplementary Table 5**. The overlap in the variants between the present study of Kerala, other Indian, and global datasets is summarized in **Supplementary Figure 1**. Out of the 4 novel variants, 1 variant in the S gene, 25281G>A, was a personal variant and was not shared by any other isolate. The remaining three novel variants were shared variants and were present in different genes (Orf1b, Orf7a and S).

**Table 1:** Summary of novel genetic variants and their frequencies in the dataset

| Variant | Gene | Variant type | Classification | Amino Acid Change | Domain/Functional Region | Frequency (Kerala N = 179) |
| --- | --- | --- | --- | --- | --- | --- |
| **21486T>C** | ORF1b | Coding mutation | Synonymous SNV | S2673S | NA | 4(2.235%) |
| **23719T>C** | S | Coding mutation | Synonymous SNV | T719T | NA | 2(1.117%) |
| **25281G>A** | S | Coding mutation | Non-Synonymous SNV | C1240Y | NA | 1(0.559%) |
| **27643C>A** | ORF7a | Coding mutation | Non-Synonymous SNV | P84T | NA | 5(2.793%) |

**Figure 3.**
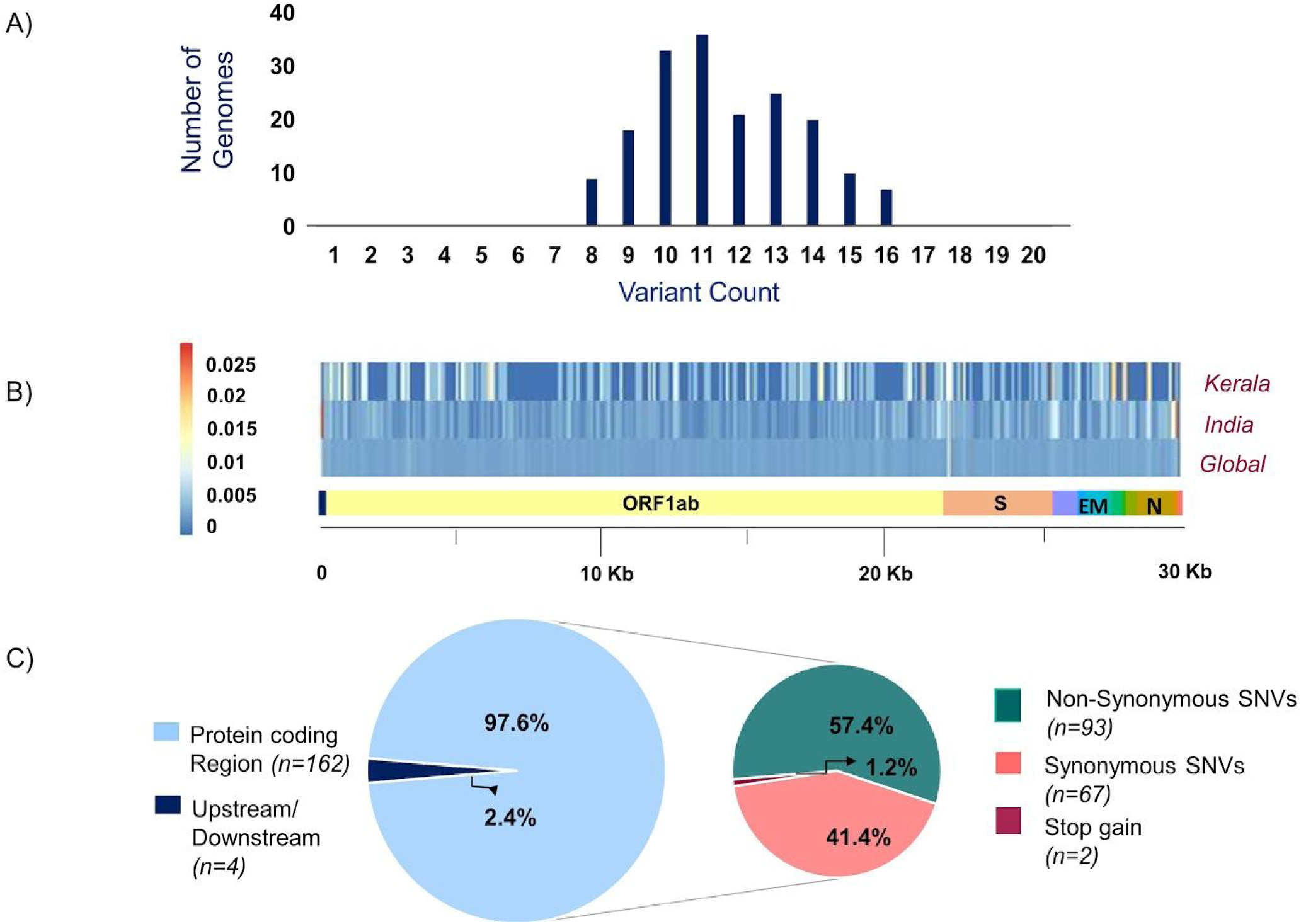
(A) Distribution of variants across genomes used in the study (B) Comparison of the proportion of the variants represented with their allele frequency across the SARS-CoV-2 genome in datasets includes Kerala (present study), India, and Global (C) Distribution of the genetic context of variants and their functional classification.

### Genomic context and classification of the variants

Of the total 166 high-quality unique variants, 162 variants were located in the protein-coding regions while 4 variants mapped to either downstream or upstream regions. Of the total variants in protein-coding regions, 93 variants were non-synonymous, 67 were synonymous, and 2 variants resulted in stopgain mutation. These two stopgain variants were found in ORF3a (26113:G>T) and ORF8 (28028:G>A) genes and were present in one individual each. The annotation of the variants based on the location and consequence is represented in **Figure 3**.

### Phylogenetic analysis

The phylogenetic tree was constructed using the genome Wuhan/WH01 (EPI_ISL_406798) as root and 2366 genomes from India which met the inclusion criteria (Ns < 5%, no missing/ambiguous date of sample collection) including 113 genomes sequenced in this study. All 113 genomes from this study were found to cluster under the globally predominant clade A2a (GISAID clade G and GH). In contrast, one of the previous genomes available from Kerala (EPI_ISL_413523, submitted by National Institute of Virology, Pune, India), which is also one of the first SARS-CoV-2 genomes sequenced in India, belongs to the clade B [13].

The dominant lineage assigned by PANGOLIN for the 113 genomes was found to be B.1 (n=110), while 3 genomes were assigned the lineage B.1.113. The phylogenetic map of the dataset of Indian genomes and the distribution of lineages in the 113 genomes from Kerala are summarized in **Figure 4**.

**Figure 4.**
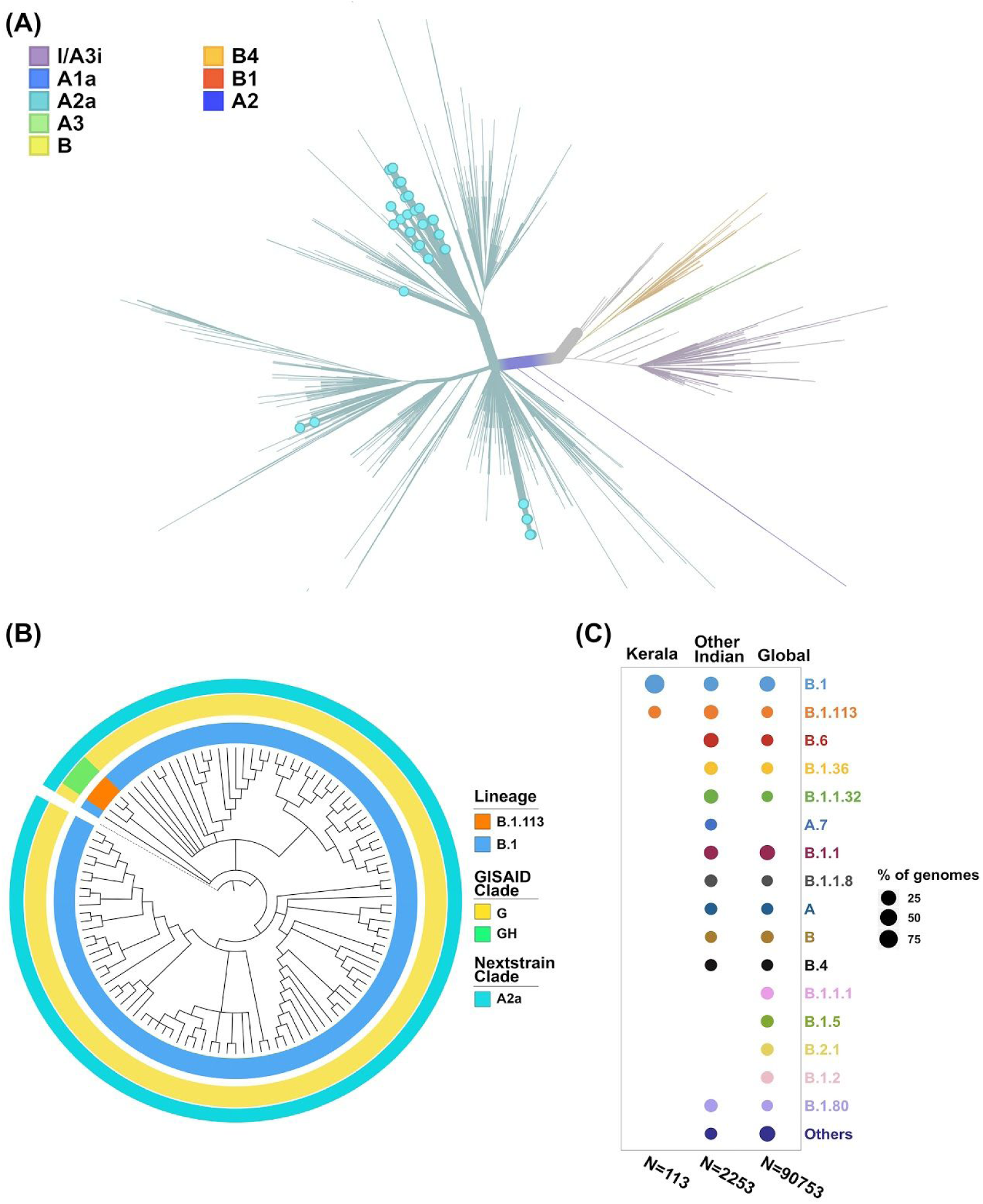
(A) Phylogenetic map of the 113 genomes sequenced from Kerala (highlighted by blue dots) with respect to the other genomes from India (B) Distribution of the clades and lineages in Kerala. All genomes clustered under the clade A2a (GISAID clade G and GH) while the dominant lineage was B.1. (C) Lineage distribution in Kerala compared to the distribution across India and global populations.

### Haplotype Analysis

Haplotype analysis was done using a dataset of 850 SARS-CoV-2 genomes from India (including 113 genomes from Kerala) that fell under clade A2a in the phylogenetic tree and clustered close to the 113 genomes from Kerala. Among the 850 genomes, there were 592 variable sites and 400 unique haplotypes **Supplementary Table 6**. The haplotype network as generated by POPART shows that a few haplogroups contributed to a majority of the isolates. Three major haplogroups contributed to 94.6% of the isolates from Kerala. The major haplogroup (K1) encompassed 40 genomes from Kerala (35.4%). The network suggests that the cluster K1 had a potential ancestor from the state of Maharashtra before possible introduction and dissemination in Kerala. A variant 16726C>T was observed to be common between the 40 genomes as well as the 3 genomes from Maharashtra belonging to the ancestral haplotype. The K1 cluster also included 4 genomes from Kerala which were found to be in a polytomy in the phylogenetic tree. Close follow-up of the cases suggests a local outbreak which contributed to the polytomy. The second haplogroup (K2) encompasses 42 genomes (37.1%) from Kerala and shares 27 genomes from Odisha. In addition, 5 genomes from Kerala in this group also constitute a polytomy. The third group (K3) encompasses 25 genomes (22.1%) from Kerala and shares 46 genomes from Karnataka. **Figure 5** summarizes the haplotype network of the A2a clade genomes.

**Figure 5.**
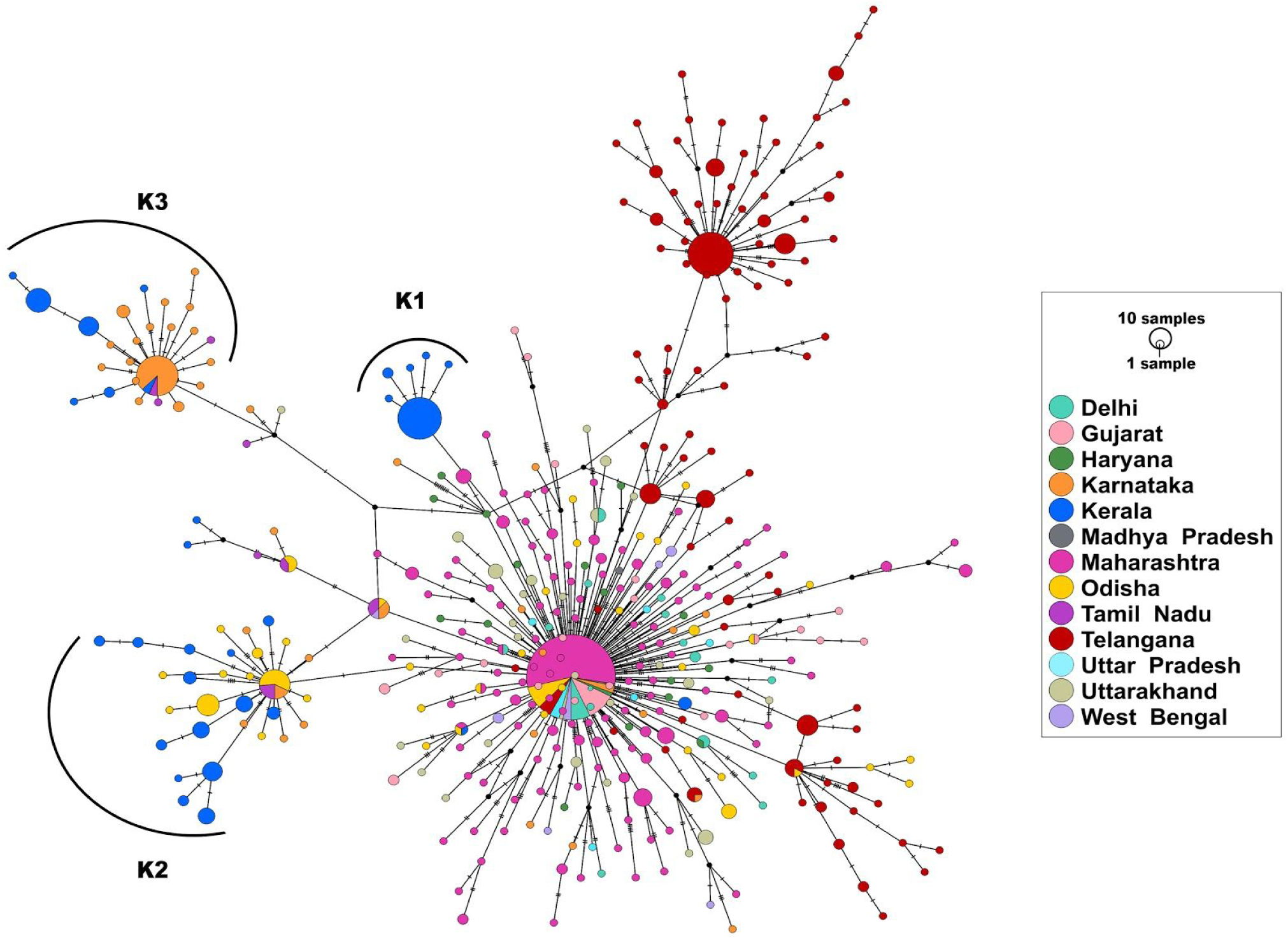
Haplotype network of 850 genomes of Indian isolates of SARS-CoV-2 belonging to the A2a clade. The 3 major haplogroups encompassing the genomes from Kerala are designated as K1, K2, and K3.

To understand the times of introduction, tMRCA was computed for the 3 distinct haplogroups. The median tMRCA were 14 July 2020 (95% highest posterior density interval [HPD] 11 May - 22 July), 20 March 2020 (95% HPD 12 Feb - 16 May), and 6 April 2020 (95% HPD 3 March - 27 May) for the three major haplogroups K1, K2, and K3 respectively. Taken together, the analysis suggests that the majority of the SARS-COV-2 isolates are outcomes of limited introductions early in the epidemic followed by local circulation.

### Functional consequences of the variants

Annotating the variants for their functional consequences using custom annotation datasets, revealed a total of 42 genetic variants that were predicted as deleterious by SIFT [37]. The filtered variants were found to span 13 unique protein domains as per UNIPROT [38] annotations. We found 15 and 120 genetic variants that mapped back to potential B and T cell epitopes from the Immune Epitope Database (IEDB) [39] respectively. In addition, 5 variants were found to span predicted error-prone sites including sequencing error sites, homoplasic positions, and hypermutable sites. Functional annotation details of all the filtered variants are summarised in **Supplementary Table 7**.

### Variants in diagnostic primer/probe binding sites in the genome

We also explored whether the variants mapped to the RT-PCR primers and probes sites. On mapping the genetic variants with the curated primers and probes, we found 5 unique variants at 5 unique primer or probes binding sites. A total of four unique variants had allele frequency > 1% at 4 unique primer binding sites. The majority of the variants i.e. 4 lies in the primer binding sites in *ORF1b, S, E*, and *N* with an allele frequency of 0.559%, 4.469%, 1.117%, and 3.352% in the Kerala isolate genomes respectively. While a variant 28899:G>T mapped to the 2019-nCoV-NFP, which is a part of China Centers for Disease Control and Prevention (CDC) primer set with a frequency of 1.117%, the Tm differed in the mutated sequence by the unit of ± 2 in comparison to the wild type sequence.

Summary of novel variants and diagnostic primer/probe spanning variants are compiled in **Table 1** and **Table 2** respectively. Details on the read count and depth of coverage of these variants are systematically documented in **Supplementary Table 8**.**a and 8**.**b**.

**Table 2:**
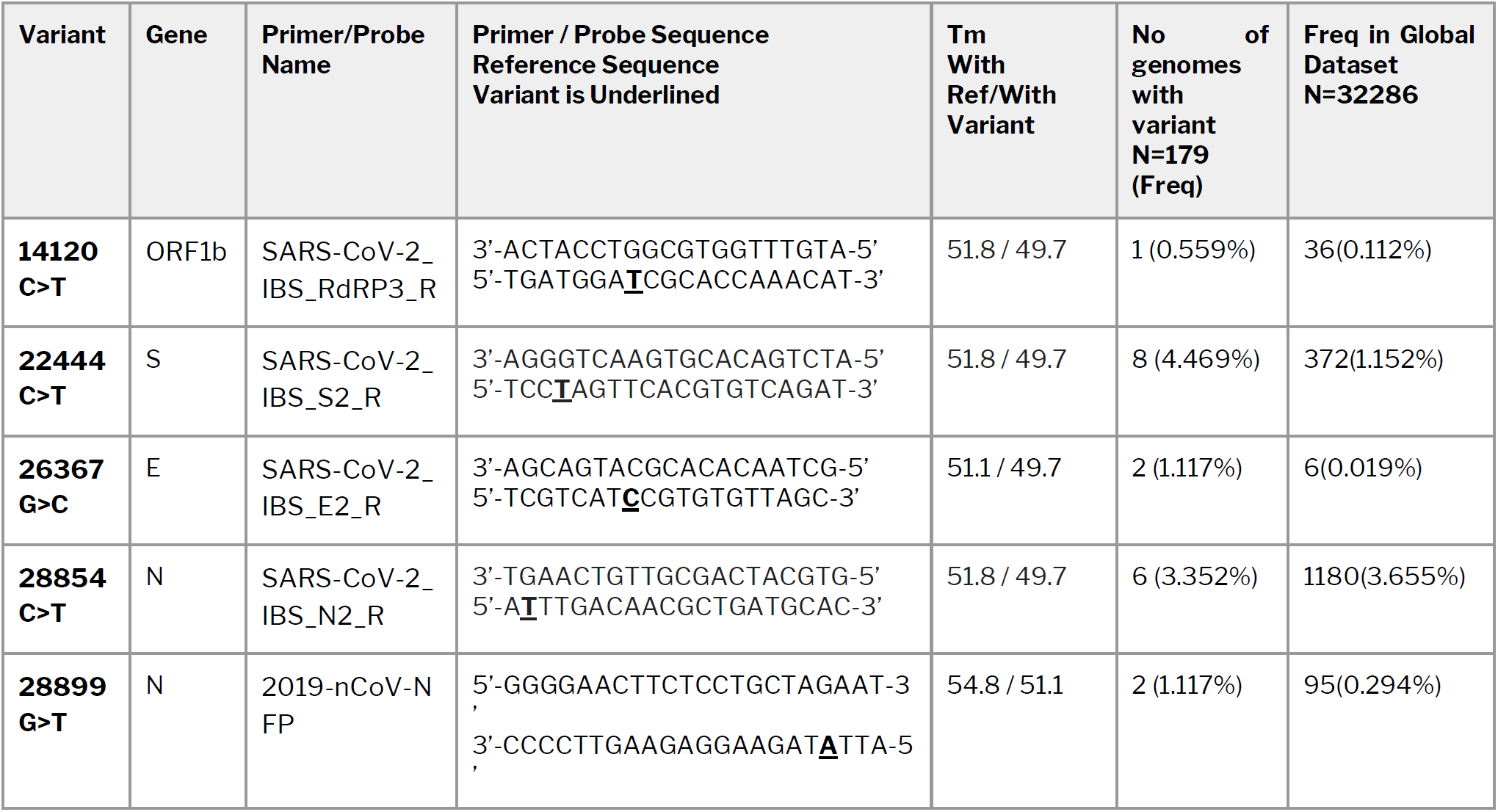
Summary of variants mapping to diagnostic RT-PCR primers or probes

### Variants associated with potential increased infectivity or attenuation of the virus in experimental settings

With the view of identifying potential functionally relevant variants, we overlapped the variants obtained from the present study with a manually curated compilation of functionally relevant SARS-CoV-2 variants. Our analysis identified 2 variants in the S gene which were reported to be associated with increased infectivity. L5F, a variation co-occurring with D614G was earlier demonstrated to possess increased infectivity [8,40] using cell line studies. In our study, 23403A>G (D614G) and 21575C>T (L5F) mutations were observed at frequencies of 99.44% and 15.64% respectively in the genomes. The combination of these variations was found to occur at a higher frequency in genomes from Kerala.

## DISCUSSION AND CONCLUSIONS

Within a small time frame, SARS-CoV-2 has spread from Wuhan (Hubei province, China) to countries across the world affecting over 26 million individuals [h ttps://covid19.who.int/]. The virus evolves by accumulating variants at an almost constant rate of 1.19 to 1.31 × 10^−3^ base substitutions per site per year [2] and therefore leaves the mutational fingerprint which is widely used for tracing the spread of the virus [41]. The availability of high-throughput sequencing approaches has enabled researchers to sequence genomes as the pandemic progressed in their respective countries. A number of methods have been adopted for rapid high throughput sequencing of SARS-CoV-2 including shotgun sequencing [4], PCR amplicon, and hybridization/capture-based enrichment and sequencing [5–7].

Genome sequencing of SARS-CoV-2 in various countries[42] has led to to insights into the temporal and geographical spread of the virus [43], introductions, and spread of the virus through travelers [44,45], local transmission, and dynamics [46], investigating the origin of outbreaks [47], just to name a few. By virtue of its connectivity to major cities through its expatriate population, trade and tourism is uniquely poised in this pandemic. It is not surprising therefore that the first case of COVID-19 in India, early in the pandemic, was reported from the state [12,13]. The genome of the isolate suggested it originated from China [13]. The following months have seen the number of cases increase to over 80 thousand in the state with a paucity of information on the origin, spread, and dynamics of the virus [https://dashboard.kerala.gov.in/].

In this present study, we performed sequencing and analysis of SARS-CoV-2 isolates from Kerala which revealed unique patterns of the transmission. These genomes are clustered into a monophyletic group mapping to the A2a clade. The A2a clade is also marked by the D614G variant, which is suggested to confer higher infectivity to the virus in experimental in vitro settings [48,49] and is therefore thought to have emerged globally as the predominant clade [8], though the cause-effect relationship still remains speculative. Haplotype analysis suggests that three major haplogroups with distinct ancestry groups encompass the majority of the isolates. The haplotype analysis in the context of other genomes from India suggests the introductions were from inter-state travel. The prevalent haplotypes were not found in any of the global genomes, supporting this observation. This also suggests that focussed testing, tracing and quarantine of expatriates and international travelers implemented during the epidemic would have been effective in curbing the spread from international travelers. The genome clusters also suggested polytomies, suggesting a recent outbreak [17]. Close follow-up of the cluster members confirmed the potential source of the outbreak, suggesting genetic epidemiology could be used in conjunction with case follow-ups to uncover potential outbreaks and possibly connect outbreaks which are apparently not related.

This study uncovered a total of 4 novel genetic variants and 89 variants which were identified only in Kerala and not in the rest of India. The genome sequences could also uncover insights into the variants of functional relevance. One of the variants of significance is a stopgain variant (28028:G>A) in the ORF8 gene. Variants including deletions in ORF8 have been suggested to attenuate the virus [50,51]. Similar variants have also been identified in other related viruses like the SARS-CoV and MERS-CoV [9,52]. A variant 21575C>T (L5F) in the *S* gene associated with increased infectivity of the virus [40] was present in 15.64% of the genomes sequenced. Following recent reports which suggest variants in the primer/probe binding sites could impact the efficiency of RT-PCR assays [11,53,54], we explored whether any of the variants in the present study mapped to the primer/probe binding sites. We identified 5 unique variants in 5 unique binding sites. The maximum number of variants were the primer set published by Won et al. [55] spanning multiple genes, apart from the 2019-nCoV-NFP GGGGAACTTCTCCTGCTAGAAT binding sites in the N gene [https://www.who.int/docs/default-source/coronaviruse/whoinhouseassays.pdf?sfvrsn=de3a76aa_2]. The latter is part of the China Centers for Disease Control and Prevention (CDC) protocol with variants in 1.117% in genomes from Kerala. We have earlier reported variants in this primer site in 39.5% of the genomes from India [11] and 18.8% [56] of global genomes. This information would be potentially valuable for laboratories in selecting reagents for screening and diagnosis.

The study has two caveats; first is that the samples were collected from a single major tertiary care center in North Kerala. However, the center caters to a large population and region and has close proximity to an international airport. Secondly, the sampling was limited to a short period of time, thus enabling only a cross-sectional view of the epidemic and precluding an accurate and temporal view of the dynamics of the epidemic in the state. Nevertheless, this provides a unique opportunity to create a snapshot of the epidemic in time and space. Notwithstanding the limitations, this is the first and most comprehensive overview of the genetic epidemiology of SARS-CoV-2 in the state of Kerala. While providing insights into the epidemiology of the epidemic, the study also enabled tracing outbreaks thereby highlighting the utility of genome sequencing as an adjunct to high-throughput screening and testing. It has not escaped our mind that scalable technologies that can combine both the approaches [7] could potentially find a place in understanding epidemics better.

## ACKNOWLEDGEMENTS

Authors thank Anjali Bajaj for editorial assistance. AJ, MD, BJ acknowledge research fellowships from CSIR. DS acknowledges a research fellowship from Intel. Authors acknowledge funding for the work from the Council of Scientific and Industrial Research (CSIR), India through grants CODEST and MLP2005. The funders had no role in the design of experiment, analysis, or decision to publish.

## SUPPLEMENTARY DATA

**Supplementary Table 1:** GISAID acknowledgment table for global genomes used in the study

**Supplementary Table 2:** GISAID acknowledgment table for genomes from India considered for phylogenetic analysis

**Supplementary Table 3:** Data summary of the samples sequenced by COVIDSeq Protocol and processed using a custom pipeline.

**Supplementary Table 4**. Summary of quality details of the variants identified in the study

**Supplementary Table 5**. Compilation of genetic variants identified for the first time in Indian genomes.

**Supplementary Table 6**. Unique haplotypes for the dataset of 850 genomes belonging to clade A2a.

**Supplementary Table 7**. Summary of functional annotation of unique genetic variants identified in the study.

**Supplementary Table 8**.**a** Tabulation of read alignment statistics for variants spanning Primer/Probe sites

**Supplementary Table 8**.**b**. Tabulation of read alignment statistics for Novel variants

**Supplementary Figure 1**. Distribution of genetic variants across datasets

