## Supplementary Table 3 for "Initial insights into the genetic epidemiology of SARS-CoV-2 isolates from Kerala suggest local spread from limited introductions"

| CS Ids | Total Reads | Trimmed Reads q30-30bp | Hisat2 Human % | Hisat2 Covidall % | Hisat2 Unmapped Covid % | BasePairescovered | X coverage | Genome coverage | # bases with 0 coverage | # bases with >0 coverage | # bases with <10x coverage | # bases with <100x coverage | BCFtools Variant Count | VARSCAN Variant Count | N's |
|---|---|---|---|---|---|---|---|---|---|---|---|---|---|---|---|
| CS1896 | 6011005 | 5696623 | 20.20% | 76.75% | 96.18% | 155440637 | 5198.161957 | 99.84616928 | 46 | 29857 | 148 | 2133 | 10 | 10 | 799 |
| CS1897 | 3951208 | 3755716 | 1.22% | 97.60% | 98.80% | 130290961 | 4357.120055 | 99.84282513 | 47 | 29856 | 58 | 102 | 11 | 10 | 15 |
| CS1898 | 6749650 | 6409736 | 4.10% | 94.47% | 98.51% | 215286007 | 7199.478547 | 99.97324683 | 8 | 29895 | 57 | 250 | 9 | 8 | 54 |
| CS1899 | 6045747 | 5756524 | 8.00% | 90.50% | 98.37% | 185230488 | 6194.378089 | 99.82944855 | 51 | 29852 | 58 | 1359 | 14 | 13 | 272 |
| CS1900 | 4188624 | 3959355 | 34.80% | 58.19% | 89.25% | 81925310 | 2739.702037 | 99.76256563 | 71 | 29832 | 393 | 2630 | 14 | 13 | 1419 |
| CS1901 | 8009923 | 7308698 | 5.78% | 92.52% | 98.19% | 240141940 | 8030.697254 | 99.91974049 | 24 | 29879 | 71 | 300 | 14 | 14 | 66 |
| CS1902 | 8737569 | 8301051 | 5.61% | 92.98% | 98.51% | 274362091 | 9175.06909 | 99.83613684 | 49 | 29854 | 58 | 373 | 14 | 13 | 18 |
| CS1903 | 12831321 | 12167248 | 0.72% | 98.44% | 99.16% | 425773231 | 14238.47878 | 99.88295489 | 35 | 29868 | 52 | 100 | 17 | 16 | 17 |
| CS1068 | 5353173 | 5053420 | 32.68% | 63.29% | 94.02% | 113711713 | 3802.685784 | 99.64217637 | 107 | 29796 | 929 | 3364 | 12 | 9 | 1023 |
| CS1069 | 5084361 | 4760062 | 65.69% | 9.43% | 27.47% | 15952745 | 533.4830953 | 85.72384042 | 4269 | 25634 | 7675 | 17118 | 11 | 9 | 7210 |
| CS1804 | 8836918 | 7607173 | 4.16% | 94.59% | 98.70% | 255217334 | 8534.840451 | 99.88295489 | 35 | 29868 | 51 | 75 | 15 | 16 | 122 |
| CS1070 | 5676866 | 5292439 | 9.64% | 88.77% | 98.24% | 166968026 | 5583.654683 | 99.91974049 | 24 | 29879 | 393 | 1968 | 8 | 8 | 683 |
| CS1071 | 6548817 | 5745212 | 1.76% | 8.50% | 8.65% | 17327554 | 579.4587165 | 88.07811925 | 3565 | 26338 | 6539 | 13378 | 7 | 6 | 6028 |
| CS1072 | 4148686 | 3907847 | 1.99% | 95.94% | 97.88% | 133299540 | 4457.731331 | 99.36795639 | 189 | 29714 | 938 | 4707 | 8 | 8 | 1301 |
| CS1073 | 4645896 | 4369665 | 1.88% | 97.30% | 99.17% | 151167324 | 5055.256128 | 99.81607197 | 55 | 29848 | 92 | 1671 | 11 | 10 | 367 |
| CS1074 | 4855750 | 4584089 | 2.91% | 96.12% | 98.99% | 156665045 | 5239.107949 | 99.86623416 | 40 | 29863 | 308 | 2478 | 10 | 9 | 905 |
| CS1075 | 5457552 | 5142926 | 61.40% | 31.30% | 81.11% | 57243911 | 1914.320001 | 91.14135705 | 2649 | 27254 | 4429 | 8295 | 13 | 9 | 4484 |
| CS1076 | 6331330 | 5990434 | 2.10% | 96.79% | 98.87% | 206117902 | 6892.883724 | 99.86289001 | 41 | 29862 | 52 | 80 | 17 | 16 | 15 |
| CS1077 | 5627062 | 4874173 | 0.69% | 98.48% | 99.17% | 170276753 | 5694.303347 | 99.84616928 | 46 | 29857 | 53 | 93 | 16 | 16 | 202 |
| CS1078 | 5931493 | 5597265 | 10.04% | 87.76% | 97.55% | 174629832 | 5839.876668 | 99.90970806 | 27 | 29876 | 83 | 819 | 15 | 14 | 340 |
| CS1079 | 7635416 | 7187029 | 2.12% | 32.96% | 33.68% | 84226986 | 2816.673444 | 99.82276026 | 53 | 29850 | 161 | 1457 | 11 | 10 | 362 |
| CS1805 | 8008259 | 7063135 | 0.13% | 99.11% | 99.23% | 248479526 | 8309.518309 | 99.9565261 | 13 | 29890 | 51 | 164 | 16 | 14 | 39 |
| CS1080 | 4946581 | 4665656 | 8.21% | 90.68% | 98.79% | 150440224 | 5030.940842 | 99.93311708 | 20 | 29883 | 343 | 2216 | 9 | 9 | 807 |
| CS1081 | 5004390 | 4720294 | 12.33% | 86.11% | 98.22% | 144536669 | 4833.517339 | 99.74918904 | 75 | 29828 | 1472 | 5217 | 9 | 9 | 1763 |
| CS1082 | 5716922 | 5345590 | 6.38% | 92.00% | 98.27% | 174788616 | 5845.186637 | 99.8261044 | 52 | 29851 | 88 | 1110 | 10 | 10 | 276 |
| CS1083 | 5144846 | 4640554 | 3.76% | 95.23% | 98.95% | 156947713 | 5248.56078 | 99.7893188 | 63 | 29840 | 763 | 4066 | 9 | 9 | 877 |
| CS1084 | 5853921 | 5531509 | 9.99% | 87.20% | 96.88% | 171518037 | 5735.813698 | 99.81941611 | 54 | 29849 | 201 | 2918 | 10 | 9 | 759 |
| CS1085 | 8326835 | 7794762 | 0.04% | 25.15% | 25.16% | 69700927 | 2330.900813 | 99.75922148 | 72 | 29831 | 927 | 3296 | 12 | 11 | 934 |
| CS1086 | 5352428 | 4932432 | 8.52% | 90.26% | 98.66% | 158161320 | 5289.145571 | 99.87961074 | 36 | 29867 | 64 | 1198 | 14 | 13 | 222 |
| CS1087 | 5533961 | 5222671 | 27.32% | 70.04% | 96.37% | 130056922 | 4349.293449 | 99.52178711 | 143 | 29760 | 914 | 2743 | 11 | 11 | 1228 |
| CS1088 | 6495097 | 6098081 | 7.53% | 18.03% | 19.50% | 39085379 | 1307.072167 | 99.12383373 | 262 | 29641 | 4024 | 7596 | 12 | 9 | 4069 |
| CS1089 | 6069624 | 5730152 | 5.90% | 91.93% | 97.69% | 187327751 | 6264.513627 | 99.83613684 | 49 | 29854 | 184 | 2398 | 11 | 10 | 1097 |
| CS1806 | 11267752 | 10211194 | 2.84% | 96.27% | 99.09% | 349056913 | 11672.97305 | 99.84951343 | 45 | 29858 | 54 | 93 | 15 | 14 | 26 |
| CS1090 | 6455867 | 6071875 | 1.20% | 97.71% | 98.90% | 210883755 | 7052.26081 | 99.85620172 | 43 | 29860 | 53 | 81 | 11 | 11 | 152 |
| CS1091 | 5716735 | 5381654 | 9.43% | 88.46% | 97.67% | 169215131 | 5658.801157 | 99.95318195 | 14 | 29889 | 59 | 514 | 12 | 12 | 226 |
| CS1092 | 5996865 | 5562880 | 2.32% | 96.77% | 99.06% | 191303763 | 6397.477277 | 99.85954586 | 42 | 29861 | 72 | 213 | 13 | 13 | 282 |
| CS1093 | 6437354 | 6046923 | 0.50% | 98.74% | 99.24% | 212236427 | 7097.496138 | 99.88295489 | 35 | 29868 | 52 | 195 | 11 | 11 | 233 |
| CS1094 | 6023960 | 5691668 | 0.53% | 98.78% | 99.31% | 199869839 | 6683.939371 | 99.85285757 | 44 | 29859 | 57 | 90 | 16 | 15 | 110 |
| CS1095 | 6228033 | 5877373 | 3.63% | 95.36% | 98.95% | 199264501 | 6663.695984 | 99.88964318 | 33 | 29870 | 76 | 1500 | 10 | 9 | 298 |
| CS1096 | 5517511 | 5200528 | 1.80% | 97.43% | 99.21% | 180128186 | 6023.749657 | 99.85285757 | 44 | 29859 | 56 | 1062 | 10 | 9 | 274 |
| CS1097 | 4627636 | 4372053 | 14.48% | 82.68% | 96.68% | 128514775 | 4297.7218 | 99.97324683 | 8 | 29895 | 127 | 2026 | 10 | 10 | 390 |
| CS1098 | 5736631 | 5403213 | 31.36% | 66.11% | 96.32% | 127000297 | 4247.075444 | 99.61207906 | 116 | 29787 | 1126 | 4086 | 11 | 11 | 1697 |
| CS1099 | 5940682 | 5602790 | 23.12% | 74.29% | 96.63% | 147982878 | 4948.763602 | 99.84282513 | 47 | 29856 | 567 | 3202 | 10 | 10 | 879 |
| CS1807 | 6423970 | 5970262 | 1.10% | 98.06% | 99.15% | 208018897 | 6956.455774 | 99.85285757 | 44 | 29859 | 51 | 105 | 15 | 15 | 267 |
| CS1100 | 6021015 | 5595798 | 12.49% | 85.63% | 97.86% | 170293566 | 5694.865599 | 99.71240344 | 86 | 29817 | 189 | 4125 | 11 | 10 | 1144 |
| CS1101 | 5032264 | 4750842 | 6.50% | 92.64% | 99.07% | 156481436 | 5232.967796 | 99.7458449 | 76 | 29827 | 937 | 4389 | 13 | 11 | 1034 |
| CS1102 | 5361918 | 5067310 | 2.92% | 95.85% | 98.73% | 172693787 | 5775.132495 | 99.92308464 | 23 | 29880 | 94 | 1463 | 12 | 12 | 327 |
| CS1103 | 5945123 | 5619237 | 0.59% | 98.71% | 99.30% | 197216753 | 6595.216299 | 99.84616928 | 46 | 29857 | 75 | 350 | 10 | 10 | 260 |
| CS1104 | 6361163 | 5992041 | 1.45% | 97.59% | 99.02% | 207870336 | 6951.487677 | 99.80269538 | 59 | 29844 | 143 | 861 | 12 | 12 | 378 |
| CS1105 | 5096896 | 4804246 | 1.78% | 97.10% | 98.85% | 165852528 | 5546.350801 | 99.82944855 | 51 | 29852 | 65 | 1460 | 11 | 11 | 348 |
| CS1106 | 5781113 | 5448832 | 0.87% | 98.33% | 99.19% | 190494249 | 6370.405946 | 99.9130522 | 26 | 29877 | 62 | 1504 | 8 | 8 | 316 |
| CS1107 | 5396943 | 5080641 | 1.58% | 96.32% | 97.87% | 174016666 | 5819.371501 | 99.69233856 | 92 | 29811 | 493 | 4359 | 17 | 16 | 1496 |
| CS1108 | 5119471 | 4813989 | 14.20% | 83.81% | 97.68% | 143408259 | 4795.781661 | 99.92977293 | 21 | 29882 | 361 | 3224 | 14 | 13 | 727 |

| | | | | | | | | | | | | | | | |
|---|---|---|---|---|---|---|---|---|---|---|---|---|---|---|---|
| CS1109 | 5392725 | 5091620 | 13.66% | 81.39% | 94.27% | 147333203 | 4927.037521 | 99.83613684 | 49 | 29854 | 113 | 1863 | 10 | 10 | 363 |
| CS1808 | 11226700 | 10614989 | 0.95% | 98.29% | 99.23% | 370901392 | 12403.48433 | 99.81941611 | 54 | 29849 | 61 | 244 | 12 | 11 | 11 |
| CS1110 | 5443316 | 5104358 | 5.40% | 93.31% | 98.64% | 169285860 | 5661.166438 | 99.83948099 | 48 | 29855 | 64 | 820 | 9 | 9 | 226 |
| CS1111 | 5449833 | 5135172 | 16.20% | 78.41% | 93.57% | 143143368 | 4786.923319 | 99.89967562 | 30 | 29873 | 88 | 1587 | 11 | 11 | 324 |
| CS1112 | 5131904 | 4816260 | 2.46% | 96.72% | 99.15% | 165569961 | 5536.901348 | 99.73915661 | 78 | 29825 | 90 | 343 | 10 | 10 | 46 |
| CS1113 | 7036316 | 6621483 | 0.96% | 98.35% | 99.30% | 231471965 | 7740.760626 | 99.83948099 | 48 | 29855 | 61 | 258 | 9 | 9 | 16 |
| CS1114 | 4803614 | 4520552 | 14.53% | 83.96% | 98.23% | 134950864 | 4512.954018 | 99.65220881 | 104 | 29799 | 749 | 4769 | 10 | 10 | 1432 |
| CS1115 | 5196983 | 4898428 | 39.15% | 56.17% | 92.31% | 97820153 | 3271.248804 | 98.92987326 | 320 | 29583 | 2331 | 6409 | 10 | 9 | 2652 |
| CS1116 | 5771368 | 5456060 | 24.28% | 73.53% | 97.10% | 142636790 | 4769.98261 | 99.07367154 | 277 | 29626 | 1066 | 3366 | 10 | 8 | 1950 |
| CS1117 | 5397037 | 4348379 | 13.02% | 85.49% | 98.28% | 131736107 | 4405.447848 | 99.5719493 | 128 | 29775 | 885 | 1373 | 15 | 13 | 1276 |
| CS1118 | 5973371 | 5619215 | 34.66% | 62.30% | 95.33% | 124454663 | 4161.945725 | 99.32782664 | 201 | 29702 | 1300 | 3145 | 10 | 9 | 2023 |
| CS1119 | 5039736 | 4750391 | 16.93% | 81.32% | 97.89% | 137352445 | 4593.266395 | 99.60873491 | 117 | 29786 | 1488 | 4874 | 10 | 9 | 1457 |
| CS1809 | 7327292 | 6956925 | 17.21% | 79.31% | 95.80% | 196165506 | 6560.061064 | 99.89633147 | 31 | 29872 | 139 | 565 | 12 | 11 | 361 |
| CS1120 | 10145 | 6364 | 0.08% | 4.05% | 4.06% | 7542 | 0.2522154968 | 0.5216867873 | 29747 | 156 | 29843 | 29872 | #N/A | #N/A | 16042 |
| CS1121 | 5049521 | 4315914 | 2.66% | 81.48% | 83.70% | 124732984 | 4171.253185 | 99.8327927 | 50 | 29853 | 74 | 1171 | 15 | 15 | 282 |
| CS1122 | 6270434 | 5885722 | 5.07% | 93.78% | 98.78% | 196182168 | 6560.618266 | 99.85954586 | 42 | 29861 | 50 | 367 | 16 | 15 | 269 |
| CS1123 | 5133354 | 4843774 | 0.97% | 98.37% | 99.33% | 169405164 | 5665.156138 | 99.81607197 | 55 | 29848 | 88 | 987 | 15 | 14 | 295 |
| CS1124 | 6356455 | 5854244 | 2.14% | 96.93% | 99.04% | 201569858 | 6740.790489 | 99.87961074 | 36 | 29867 | 63 | 226 | 13 | 13 | 292 |
| CS1125 | 5542178 | 5161203 | 3.70% | 95.22% | 98.88% | 174672459 | 5841.302177 | 99.83948099 | 48 | 29855 | 99 | 1461 | 14 | 14 | 298 |
| CS1126 | 5035121 | 4655388 | 3.65% | 94.52% | 98.10% | 156346481 | 5228.454704 | 99.91639635 | 25 | 29878 | 86 | 1038 | 13 | 13 | 336 |
| CS1127 | 5709455 | 4769793 | 2.69% | 95.92% | 98.58% | 162227019 | 5425.108484 | 99.84616928 | 46 | 29857 | 73 | 879 | 13 | 13 | 233 |
| CS1128 | 6219490 | 5862076 | 21.45% | 76.52% | 97.42% | 159479982 | 5333.243554 | 99.8261044 | 52 | 29851 | 478 | 2348 | 9 | 9 | 815 |
| CS1129 | 4102886 | 3879629 | 1.97% | 97.26% | 99.21% | 134115116 | 4485.005384 | 99.85285757 | 44 | 29859 | 60 | 103 | 10 | 10 | 55 |
| CS1810 | 10392691 | 9838901 | 1.65% | 97.52% | 99.16% | 341062908 | 11405.64184 | 99.8261044 | 52 | 29851 | 65 | 258 | 11 | 11 | 17 |
| CS1130 | 5420458 | 5120076 | 5.84% | 93.11% | 98.88% | 169490605 | 5668.01341 | 99.91639635 | 25 | 29878 | 90 | 1511 | 12 | 12 | 394 |
| CS1131 | 6012270 | 5692963 | 2.63% | 96.51% | 99.12% | 195356578 | 6533.00933 | 99.84616928 | 46 | 29857 | 57 | 693 | 10 | 9 | 278 |
| CS1132 | 4857107 | 4574339 | 23.31% | 74.38% | 96.99% | 120968354 | 4045.358459 | 99.87961074 | 36 | 29867 | 231 | 611 | 9 | 10 | 1010 |
| CS1133 | 5128420 | 4718889 | 17.75% | 80.84% | 98.28% | 135554087 | 4533.126676 | 99.69902685 | 90 | 29813 | 521 | 5670 | 9 | 10 | 1594 |
| CS1134 | 7377548 | 6971926 | 5.18% | 93.95% | 99.09% | 232902756 | 7788.608367 | 99.85620172 | 43 | 29860 | 85 | 857 | 10 | 10 | 307 |
| CS1135 | 6159498 | 5791815 | 16.01% | 38.43% | 45.75% | 79148470 | 2646.840451 | 95.17439722 | 1443 | 28460 | 4522 | 7427 | 10 | 11 | 4180 |
| CS1136 | 5949510 | 5595842 | 10.16% | 87.92% | 97.87% | 174881623 | 5848.296927 | 99.92642879 | 22 | 29881 | 62 | 1077 | 12 | 13 | 341 |
| CS1137 | 6140108 | 5798311 | 3.54% | 95.64% | 99.15% | 197153667 | 6593.106611 | 99.85285757 | 44 | 29859 | 75 | 189 | 15 | 14 | 280 |
| CS1138 | 5636748 | 5322714 | 9.19% | 88.45% | 97.40% | 167377449 | 5597.346387 | 99.93980537 | 18 | 29885 | 75 | 523 | 15 | 15 | 319 |
| CS1139 | 6614597 | 6164809 | 47.60% | 33.00% | 62.98% | 72326653 | 2418.708926 | 97.1474434 | 853 | 29050 | 3333 | 6929 | 15 | 13 | 3173 |
| CS1811 | 8290150 | 7852724 | 3.80% | 95.09% | 98.85% | 265462483 | 8877.453199 | 99.84282513 | 47 | 29856 | 54 | 102 | 14 | 14 | 139 |
| CS1140 | 6151367 | 5809823 | 0.36% | 98.49% | 98.85% | 203412925 | 6802.425342 | 99.89298733 | 32 | 29871 | 51 | 322 | 12 | 12 | 279 |
| CS1141 | 6046396 | 5705075 | 11.06% | 87.52% | 98.41% | 177519204 | 5936.501488 | 99.7458449 | 76 | 29827 | 331 | 1316 | 11 | 10 | 504 |
| CS1142 | 6405889 | 6018588 | 8.14% | 90.71% | 98.76% | 194080646 | 6490.3403 | 99.83948099 | 48 | 29855 | 77 | 1350 | 10 | 10 | 297 |
| CS1143 | 4084746 | 3845510 | 5.62% | 93.30% | 98.85% | 127548315 | 4265.401966 | 99.81607197 | 55 | 29848 | 76 | 519 | 10 | 10 | 278 |
| CS1144 | 8004094 | 7567378 | 3.32% | 95.69% | 98.97% | 257382610 | 8607.250443 | 99.8327927 | 50 | 29853 | 58 | 365 | 10 | 10 | 187 |
| CS1145 | 6768381 | 6400431 | 3.30% | 95.78% | 99.05% | 217911639 | 7287.283517 | 99.94649366 | 16 | 29887 | 56 | 113 | 17 | 16 | 90 |
| CS1146 | 4999181 | 4510776 | 2.23% | 93.66% | 95.79% | 149992273 | 5015.960706 | 99.96990269 | 9 | 29894 | 59 | 1309 | 11 | 11 | 272 |
| CS1147 | 5062479 | 4692223 | 8.65% | 61.07% | 66.86% | 101845328 | 3405.856536 | 99.87292245 | 38 | 29865 | 1075 | 3190 | 11 | 11 | 954 |
| CS1148 | 5372254 | 5027203 | 4.53% | 94.61% | 99.10% | 169061261 | 5653.65552 | 99.8695783 | 39 | 29864 | 86 | 968 | 10 | 10 | 312 |
| CS1149 | 4968156 | 4698955 | 9.66% | 89.13% | 98.66% | 148900185 | 4979.439688 | 99.8261044 | 52 | 29851 | 336 | 1971 | 11 | 11 | 712 |
| CS1812 | 7551226 | 6898516 | 1.90% | 97.26% | 99.14% | 238293270 | 7968.875029 | 99.85620172 | 43 | 29860 | 57 | 429 | 15 | 14 | 211 |
| CS1150 | 5890375 | 5495519 | 14.71% | 83.85% | 98.32% | 163773974 | 5476.840919 | 99.7893188 | 63 | 29840 | 737 | 4036 | 12 | 10 | 721 |
| CS1151 | 7104529 | 6667738 | 1.28% | 98.03% | 99.30% | 232305403 | 7768.63201 | 99.84282513 | 47 | 29856 | 55 | 107 | 11 | 11 | 20 |
| CS1152 | 5522232 | 5199706 | 6.04% | 92.54% | 98.49% | 171048639 | 5720.116343 | 99.88629903 | 34 | 29869 | 91 | 939 | 11 | 11 | 304 |
| CS1153 | 6480588 | 6021794 | 1.54% | 97.77% | 99.30% | 209213620 | 6996.409056 | 99.82944855 | 51 | 29852 | 89 | 1084 | 11 | 11 | 295 |
| CS1154 | 5849220 | 5516893 | 7.92% | 90.97% | 98.79% | 178418451 | 5966.573621 | 99.75922148 | 72 | 29831 | 138 | 1317 | 12 | 12 | 320 |
| CS1155 | 5884203 | 5550712 | 1.06% | 98.30% | 99.35% | 193983744 | 6487.099756 | 99.81272782 | 56 | 29847 | 88 | 705 | 12 | 12 | 265 |
| CS1156 | 6099940 | 5317785 | 2.98% | 96.02% | 98.98% | 181183179 | 6059.030164 | 99.95318195 | 14 | 29889 | 75 | 367 | 11 | 11 | 73 |
| CS1157 | 5245255 | 4943017 | 18.57% | 79.59% | 97.74% | 139871507 | 4677.507508 | 99.80269538 | 59 | 29844 | 457 | 2122 | 11 | 10 | 767 |
| CS1158 | 5648901 | 4852302 | 21.09% | 75.68% | 95.91% | 130284376 | 4356.899843 | 99.81272782 | 56 | 29847 | 1191 | 4543 | 12 | 11 | 1173 |

| | | | | | | | | | | | | | | | |
|---|---|---|---|---|---|---|---|---|---|---|---|---|---|---|---|
| CS1159 | 5287524 | 4990529 | 0.81% | 98.29% | 99.09% | 174370230 | 5831.195198 | 99.8327927 | 50 | 29853 | 68 | 1455 | 11 | 11 | 296 |
| CS1813 | 8206194 | 7756400 | 3.68% | 95.09% | 98.73% | 262225751 | 8769.212153 | 99.72578002 | 82 | 29821 | 145 | 1052 | 13 | 13 | 1464 |
| CS1795 | 14245366 | 13509456 | 0.58% | 98.34% | 98.91% | 472165213 | 15789.89443 | 99.85285757 | 44 | 29859 | 50 | 57 | 10 | 10 | 11 |
| CS1160 | 6548309 | 6172648 | 1.47% | 97.54% | 98.99% | 214005381 | 7156.652543 | 99.96990269 | 9 | 29894 | 64 | 153 | 12 | 11 | 253 |
| CS1814 | 9563688 | 9028069 | 1.13% | 98.19% | 99.31% | 315070739 | 10536.42574 | 99.8327927 | 50 | 29853 | 89 | 193 | 14 | 13 | 160 |
| CS1815 | 10019608 | 9523894 | 1.30% | 97.97% | 99.26% | 331691180 | 11092.23757 | 99.8261044 | 52 | 29851 | 83 | 99 | 14 | 14 | 75 |
| CS1816 | 9664860 | 9055485 | 2.31% | 96.83% | 99.12% | 311597712 | 10420.28265 | 99.8695783 | 39 | 29864 | 57 | 101 | 14 | 13 | 44 |
| CS1817 | 13264607 | 12550591 | 0.35% | 98.92% | 99.26% | 441380095 | 14760.39511 | 99.94649366 | 16 | 29887 | 43 | 56 | 14 | 14 | 29 |
| CS1818 | 9879870 | 8928739 | 0.38% | 98.94% | 99.32% | 313668287 | 10489.5257 | 99.81272782 | 56 | 29847 | 88 | 107 | 12 | 12 | 37 |
| CS1819 | 9909813 | 9388142 | 0.88% | 98.30% | 99.16% | 328026414 | 10969.68244 | 99.87961074 | 36 | 29867 | 53 | 83 | 14 | 13 | 27 |
| CS1820 | 5479996 | 5096243 | 19.00% | 77.53% | 95.72% | 140371770 | 4694.237033 | 99.5719493 | 128 | 29775 | 226 | 1259 | 8 | 8 | 1808 |
| CS1821 | 13570547 | 12878688 | 1.24% | 97.90% | 99.13% | 448145526 | 14986.64101 | 99.86289001 | 41 | 29862 | 49 | 70 | 14 | 14 | 24 |
| CS1822 | 13849926 | 11640395 | 2.05% | 96.88% | 98.91% | 399920049 | 13373.91061 | 99.83948099 | 48 | 29855 | 85 | 560 | 8 | 8 | 237 |
| CS1823 | 16688459 | 15856965 | 0.63% | 98.63% | 99.25% | 555999835 | 18593.44664 | 99.81607197 | 55 | 29848 | 83 | 112 | 11 | 11 | 51 |
| CS1796 | 1482384 | 1383619 | 87.94% | 8.31% | 68.90% | 4085435 | 136.6229141 | 6.464234358 | 27970 | 1933 | 28167 | 28293 | 1 | 1 | 28200 |
| CS1824 | 16889111 | 16043578 | 1.72% | 97.29% | 98.99% | 554904623 | 18556.82116 | 99.83948099 | 48 | 29855 | 53 | 96 | 11 | 10 | 10 |
| CS1825 | 21250504 | 20094373 | 19.94% | 72.08% | 90.03% | 514940775 | 17220.3717 | 99.96655854 | 10 | 29893 | 49 | 531 | 11 | 10 | 227 |
| CS1826 | 20228344 | 17700371 | 4.87% | 93.94% | 98.75% | 589966901 | 19729.35495 | 99.95987025 | 12 | 29891 | 82 | 258 | 11 | 11 | 79 |
| CS1827 | 16424777 | 15630659 | 19.73% | 76.03% | 94.72% | 422534241 | 14130.16222 | 99.92308464 | 23 | 29880 | 217 | 641 | 11 | 11 | 209 |
| CS1828 | 9365061 | 8864461 | 1.96% | 96.88% | 98.82% | 305234065 | 10207.473 | 99.79935124 | 60 | 29843 | 208 | 255 | 10 | 10 | 156 |
| CS1829 | 9742635 | 8563813 | 0.60% | 98.53% | 99.13% | 299470293 | 10014.72404 | 99.87292245 | 38 | 29865 | 53 | 67 | 14 | 13 | 19 |
| CS1830 | 13027402 | 12348495 | 1.74% | 97.29% | 99.01% | 427035847 | 14280.7025 | 99.86289001 | 41 | 29862 | 49 | 65 | 13 | 12 | 10 |
| CS1831 | 11622127 | 11006082 | 3.40% | 94.63% | 97.97% | 370285987 | 12382.90429 | 99.96990269 | 9 | 29894 | 91 | 258 | 12 | 12 | 75 |
| CS1832 | 11566356 | 10885741 | 2.46% | 96.65% | 99.09% | 373887270 | 12503.33645 | 99.84282513 | 47 | 29856 | 59 | 214 | 12 | 11 | 104 |
| CS1833 | 11070971 | 9829766 | 16.37% | 76.16% | 91.07% | 265730674 | 8886.421897 | 99.92308464 | 23 | 29880 | 217 | 704 | 12 | 12 | 285 |
| CS1797 | 7209742 | 6674603 | 1.93% | 97.25% | 99.16% | 230586562 | 7711.151456 | 99.85285757 | 44 | 29859 | 56 | 108 | 15 | 15 | 101 |
| CS1836 | 10711499 | 10162836 | 2.37% | 96.57% | 98.91% | 348887734 | 11667.31545 | 99.84616928 | 46 | 29857 | 53 | 78 | 13 | 13 | 16 |
| CS1837 | 12965575 | 12285708 | 7.46% | 88.87% | 96.03% | 388100114 | 12978.63472 | 99.9565261 | 13 | 29890 | 30 | 258 | 11 | 12 | 30 |
| CS1838 | 14720595 | 13817560 | 13.84% | 84.06% | 97.56% | 412767884 | 13803.56098 | 99.97324683 | 8 | 29895 | 54 | 217 | 11 | 10 | 227 |
| CS1839 | 14124594 | 12516982 | 10.53% | 86.89% | 97.11% | 385964710 | 12907.22369 | 99.97993512 | 6 | 29897 | 71 | 282 | 11 | 11 | 73 |
| CS1840 | 16023748 | 15167013 | 3.45% | 94.67% | 98.05% | 510448741 | 17070.15152 | 99.96655854 | 10 | 29893 | 26 | 244 | 12 | 12 | 55 |
| CS1841 | 18245950 | 17280821 | 4.31% | 94.50% | 98.76% | 580488786 | 19412.39294 | 99.87961074 | 36 | 29867 | 50 | 85 | 9 | 9 | 68 |
| CS1842 | 22238804 | 21142118 | 5.16% | 93.60% | 98.69% | 703486750 | 23525.62452 | 99.98327927 | 5 | 29898 | 33 | 143 | 10 | 8 | 68 |
| CS1843 | 25088909 | 23842703 | 3.57% | 95.32% | 98.85% | 807762809 | 27012.76825 | 99.88964318 | 33 | 29870 | 45 | 56 | 13 | 13 | 14 |
| CS1844 | 6365048 | 6055513 | 1.96% | 97.14% | 99.08% | 209167866 | 6994.878975 | 99.8261044 | 52 | 29851 | 158 | 985 | 14 | 12 | 595 |
| CS1845 | 7386514 | 6453383 | 21.52% | 73.36% | 93.47% | 167972616 | 5617.249641 | 99.94649366 | 16 | 29887 | 74 | 1027 | 14 | 14 | 355 |
| CS1798 | 6689743 | 6340321 | 5.61% | 93.37% | 98.92% | 210450170 | 7037.761094 | 99.8261044 | 52 | 29851 | 61 | 664 | 12 | 12 | 210 |
| CS1846 | 9802962 | 9310696 | 8.71% | 89.57% | 98.12% | 296508334 | 9915.671806 | 99.97659098 | 7 | 29896 | 58 | 416 | 15 | 14 | 242 |
| CS1847 | 13310793 | 12631620 | 11.45% | 85.32% | 96.35% | 383110626 | 12811.77895 | 99.94314952 | 17 | 29886 | 76 | 391 | 12 | 11 | 71 |
| CS1848 | 13782250 | 13039460 | 18.23% | 72.86% | 89.11% | 337707649 | 11293.43708 | 99.95987025 | 12 | 29891 | 45 | 280 | 9 | 9 | 109 |
| CS1849 | 4705257 | 4462810 | 0.00% | 97.88% | 97.88% | 155335201 | 5194.636023 | 37.33070261 | 18740 | 11163 | 19554 | 19606 | 4 | 4 | 25428 |
| CS1850 | 17808366 | 16748123 | 12.08% | 86.20% | 98.04% | 513276753 | 17164.72438 | 99.63883222 | 108 | 29795 | 444 | 1716 | 9 | 9 | 679 |
| CS1851 | 17069993 | 15512013 | 12.66% | 85.59% | 97.99% | 471415879 | 15764.8356 | 99.95987025 | 12 | 29891 | 62 | 159 | 10 | 10 | 240 |
| CS1852 | 4741354 | 4484460 | 62.29% | 19.57% | 51.90% | 31210163 | 1043.71344 | 97.98347992 | 603 | 29300 | 3630 | 9455 | 11 | 10 | 3652 |
| CS1853 | 7053975 | 6674559 | 2.54% | 96.30% | 98.81% | 228445499 | 7639.551182 | 99.92977293 | 21 | 29882 | 59 | 109 | 15 | 13 | 55 |
| CS1854 | 6472320 | 5993896 | 1.28% | 97.57% | 98.83% | 207738046 | 6947.063706 | 99.84616928 | 46 | 29857 | 76 | 292 | 16 | 15 | 198 |
| CS1855 | 7352891 | 6973484 | 2.21% | 89.99% | 92.03% | 223085915 | 7460.318864 | 99.85954586 | 42 | 29861 | 61 | 474 | 14 | 13 | 225 |
| CS1799 | 7997214 | 7536700 | 3.40% | 95.68% | 99.06% | 256310386 | 8571.393706 | 99.84951343 | 45 | 29858 | 55 | 100 | 12 | 12 | 47 |
| CS1856 | 10209896 | 9654205 | 4.11% | 93.55% | 97.56% | 320966404 | 10733.58539 | 99.98662342 | 4 | 29899 | 52 | 93 | 11 | 11 | 181 |
| CS1857 | 19995846 | 18898935 | 0.20% | 11.79% | 11.81% | 79200672 | 2648.586162 | 98.78273083 | 364 | 29539 | 2143 | 5576 | 11 | 10 | 2406 |
| CS1858 | 13661481 | 12910290 | 9.35% | 88.85% | 98.01% | 407807807 | 13637.68876 | 99.81272782 | 56 | 29847 | 202 | 1836 | 10 | 10 | 832 |
| CS1859 | 13824886 | 13082875 | 4.24% | 93.47% | 97.61% | 434635235 | 14534.83714 | 99.85285757 | 44 | 29859 | 61 | 224 | 10 | 10 | 65 |
| CS1860 | 4754606 | 4443090 | 14.15% | 83.83% | 97.65% | 132358321 | 4426.255593 | 99.67896198 | 96 | 29807 | 347 | 2102 | 13 | 13 | 549 |
| CS1861 | 5801320 | 5482161 | 9.91% | 88.19% | 97.88% | 171828670 | 5746.201719 | 99.93311708 | 20 | 29883 | 202 | 1040 | 13 | 13 | 379 |
| CS1862 | 8846456 | 8351640 | 0.42% | 11.44% | 11.49% | 33968313 | 1135.950005 | 94.34170485 | 1692 | 28211 | 4575 | 7732 | 10 | 10 | 4252 |

| | | | | | | | | | | | | | | | |
|---|---|---|---|---|---|---|---|---|---|---|---|---|---|---|---|
| CS1863 | 6978541 | 6623326 | 17.23% | 80.51% | 97.27% | 189616735 | 6341.060596 | 99.81941611 | 54 | 29849 | 129 | 1587 | 11 | 11 | 494 |
| CS1864 | 9743342 | 9241040 | 2.21% | 96.18% | 98.36% | 315950650 | 10565.85125 | 99.87961074 | 36 | 29867 | 65 | 137 | 8 | 8 | 38 |
| CS1865 | 7890781 | 7507101 | 3.99% | 84.65% | 88.17% | 225959821 | 7556.426479 | 99.81941611 | 54 | 29849 | 59 | 878 | 14 | 14 | 277 |
| CS1800 | 9138561 | 8613764 | 0.83% | 98.30% | 99.13% | 300972721 | 10064.96743 | 99.84616928 | 46 | 29857 | 59 | 103 | 13 | 13 | 48 |
| CS1866 | 10739141 | 10174272 | 28.81% | 66.49% | 93.40% | 240520038 | 8043.341404 | 99.77594221 | 67 | 29836 | 249 | 1112 | 14 | 12 | 504 |
| CS1867 | 12067090 | 11440050 | 2.85% | 96.01% | 98.82% | 390429609 | 13056.53643 | 99.97324683 | 8 | 29895 | 56 | 101 | 12 | 12 | 60 |
| CS1868 | 7811122 | 7239328 | 1.67% | 97.00% | 98.65% | 249448349 | 8341.917166 | 99.87292245 | 38 | 29865 | 55 | 100 | 14 | 13 | 22 |
| CS1869 | 9222564 | 8752168 | 0.13% | 99.13% | 99.26% | 308420713 | 10314.03916 | 99.95318195 | 14 | 29889 | 53 | 83 | 14 | 13 | 43 |
| CS1870 | 8114031 | 7714832 | 1.65% | 97.39% | 99.03% | 267111956 | 8932.613985 | 99.84282513 | 47 | 29856 | 58 | 95 | 16 | 16 | 162 |
| CS1871 | 8422981 | 7999844 | 14.02% | 82.84% | 96.35% | 235632994 | 7879.911514 | 99.97659098 | 7 | 29896 | 37 | 499 | 16 | 14 | 277 |
| CS1872 | 12723229 | 12055008 | 5.51% | 92.87% | 98.29% | 397993588 | 13309.48694 | 99.97993512 | 6 | 29897 | 23 | 80 | 15 | 15 | 249 |
| CS1873 | 8915258 | 8450913 | 4.72% | 91.87% | 96.42% | 275938635 | 9227.791024 | 99.98327927 | 5 | 29898 | 56 | 163 | 15 | 15 | 77 |
| CS1874 | 9904191 | 9379599 | 24.54% | 68.35% | 90.57% | 227922632 | 7622.065746 | 99.96321439 | 11 | 29892 | 58 | 1205 | 16 | 15 | 313 |
| CS1875 | 8236175 | 7810987 | 15.99% | 81.53% | 97.04% | 226392674 | 7570.901716 | 99.81607197 | 55 | 29848 | 65 | 669 | 11 | 10 | 274 |
| CS1801 | 9030161 | 8563297 | 17.64% | 75.76% | 91.98% | 230680680 | 7714.2989 | 99.90970806 | 27 | 29876 | 197 | 921 | 11 | 11 | 305 |
| CS1876 | 4376255 | 4141672 | 20.09% | 76.59% | 95.84% | 112781465 | 3771.576932 | 99.90301976 | 29 | 29874 | 424 | 2164 | 11 | 9 | 890 |
| CS1877 | 7613296 | 7235916 | 0.56% | 98.55% | 99.11% | 253494590 | 8477.229375 | 99.8762666 | 37 | 29866 | 54 | 79 | 15 | 14 | 31 |
| CS1878 | 9312963 | 8778144 | 0.23% | 98.89% | 99.12% | 308487553 | 10316.27439 | 99.87292245 | 38 | 29865 | 51 | 65 | 15 | 14 | 29 |
| CS1879 | 17070963 | 16192229 | 0.57% | 98.73% | 99.30% | 568283166 | 19004.21918 | 99.85954586 | 42 | 29861 | 49 | 58 | 12 | 11 | 9 |
| CS1880 | 11352159 | 10738152 | 3.49% | 95.55% | 99.00% | 364738383 | 12197.38431 | 99.85285757 | 44 | 29859 | 58 | 365 | 9 | 9 | 226 |
| CS1881 | 12679877 | 12018730 | 23.01% | 74.83% | 97.20% | 319803200 | 10694.68615 | 99.75253319 | 74 | 29829 | 88 | 1067 | 9 | 9 | 619 |
| CS1882 | 12863553 | 12180097 | 91.01% | 3.13% | 34.83% | 13565201 | 453.6401364 | 18.83757483 | 24270 | 5633 | 25890 | 26280 | 5 | 2 | 25735 |
| CS1883 | 14694899 | 13968401 | 8.10% | 90.42% | 98.39% | 449006435 | 15015.43106 | 99.82944855 | 51 | 29852 | 70 | 318 | 10 | 10 | 272 |
| CS1884 | 6374062 | 6052916 | 2.08% | 96.94% | 99.01% | 208637897 | 6977.156038 | 99.82276026 | 53 | 29850 | 134 | 1581 | 11 | 11 | 537 |
| CS1885 | 6867035 | 5544600 | 10.10% | 87.20% | 96.99% | 171315991 | 5729.056984 | 99.8261044 | 52 | 29851 | 67 | 1007 | 11 | 11 | 172 |
| CS1802 | 10169376 | 9563454 | 6.21% | 90.74% | 96.75% | 308472704 | 10315.77781 | 99.92977293 | 21 | 29882 | 138 | 366 | 8 | 8 | 115 |
| CS1886 | 7407713 | 6998256 | 1.46% | 29.36% | 29.79% | 73037709 | 2442.487677 | 99.41143029 | 176 | 29727 | 1417 | 5177 | 11 | 11 | 1479 |
| CS1887 | 7484378 | 7097486 | 2.93% | 95.83% | 98.72% | 241794536 | 8085.962479 | 99.88295489 | 35 | 29868 | 63 | 224 | 11 | 11 | 29 |
| CS1888 | 9688673 | 9190783 | 2.63% | 96.26% | 98.86% | 314527406 | 10518.25589 | 99.86289001 | 41 | 29862 | 52 | 90 | 12 | 12 | 191 |
| CS1889 | 8262427 | 7126058 | 3.42% | 95.21% | 98.58% | 240670956 | 8048.388322 | 99.84282513 | 47 | 29856 | 52 | 84 | 11 | 11 | 271 |
| CS1890 | 9920499 | 9365888 | 1.72% | 96.72% | 98.42% | 321934613 | 10765.96372 | 99.88295489 | 35 | 29868 | 48 | 63 | 11 | 11 | 199 |
| CS1891 | 8739719 | 8290784 | 4.60% | 82.11% | 86.07% | 242057737 | 8094.764305 | 99.84951343 | 45 | 29858 | 59 | 360 | 9 | 8 | 339 |
| CS1892 | 5302251 | 4922068 | 2.07% | 96.94% | 98.99% | 169511948 | 5668.727151 | 99.83613684 | 49 | 29854 | 56 | 1228 | 12 | 12 | 241 |
| CS1893 | 5822983 | 5456387 | 8.25% | 90.44% | 98.58% | 175403279 | 5865.741865 | 99.95318195 | 14 | 29889 | 59 | 1215 | 13 | 12 | 365 |
| CS1894 | 4630806 | 4307798 | 6.16% | 91.97% | 98.01% | 140779644 | 4707.876935 | 99.81272782 | 56 | 29847 | 86 | 648 | 15 | 14 | 296 |
| CS1895 | 6033648 | 5138672 | 18.31% | 79.19% | 96.94% | 144359596 | 4827.59576 | 97.07387219 | 875 | 29028 | 1694 | 3176 | 17 | 16 | 2661 |
| CS1803 | 9918776 | 9402455 | 1.15% | 97.94% | 99.08% | 327320719 | 10946.08297 | 99.85954586 | 42 | 29861 | 49 | 61 | 15 | 14 | 12 |
