## Supplementary Table 4 for "Initial insights into the genetic epidemiology of SARS-CoV-2 isolates from Kerala suggest local spread from limited introductions"

| Chr | Pos | Ref | Alt | Occurrence | Average Variation Percentage |
|---|---|---|---|---|---|
| 1 | 2334 | C | T | 1 | 100 |
| 1 | 2523 | C | T | 1 | 100 |
| 1 | 2875 | G | T | 1 | 100 |
| 1 | 3604 | C | T | 1 | 100 |
| 1 | 3728 | G | A | 4 | 100 |
| 1 | 4754 | C | T | 1 | 100 |
| 1 | 5812 | C | G | 1 | 100 |
| 1 | 5833 | C | T | 1 | 100 |
| 1 | 5866 | C | T | 1 | 100 |
| 1 | 6402 | C | T | 1 | 100 |
| 1 | 8371 | G | T | 4 | 100 |
| 1 | 8782 | C | T | 1 | 100 |
| 1 | 9707 | A | G | 1 | 100 |
| 1 | 11950 | C | T | 1 | 100 |
| 1 | 13957 | C | T | 1 | 100 |
| 1 | 14120 | C | T | 1 | 100 |
| 1 | 16402 | G | T | 4 | 100 |
| 1 | 19017 | C | T | 1 | 100 |
| 1 | 20302 | G | A | 2 | 100 |
| 1 | 21017 | C | T | 2 | 100 |
| 1 | 21575 | C | T | 28 | 100 |
| 1 | 21621 | C | T | 9 | 100 |
| 1 | 21846 | C | T | 1 | 100 |
| 1 | 21952 | C | T | 1 | 100 |
| 1 | 22201 | G | T | 2 | 100 |
| 1 | 22444 | C | T | 8 | 100 |
| 1 | 22448 | C | A | 3 | 100 |
| 1 | 22468 | G | T | 1 | 100 |
| 1 | 23248 | C | T | 1 | 100 |
| 1 | 23719 | T | C | 2 | 100 |
| 1 | 24130 | C | T | 1 | 100 |
| 1 | 24933 | G | T | 1 | 100 |
| 1 | 25511 | C | T | 1 | 100 |
| 1 | 25546 | C | T | 1 | 100 |
| 1 | 26486 | T | A | 1 | 100 |
| 1 | 28024 | A | G | 1 | 100 |
| 1 | 28028 | G | A | 1 | 100 |
| 1 | 28878 | G | A | 1 | 100 |
| 1 | 28980 | G | T | 1 | 100 |
| 1 | 29614 | C | T | 1 | 100 |
| 1 | 29659 | C | T | 4 | 100 |
| 1 | 28854 | C | T | 6 | 99.9967 |
| 1 | 16726 | C | T | 47 | 99.9962 |
| 1 | 18129 | C | T | 9 | 99.9956 |
| 1 | 3653 | C | T | 9 | 99.9944 |
| 1 | 12017 | T | G | 4 | 99.9925 |
| 1 | 12325 | C | T | 6 | 99.9917 |
| 1 | 26735 | C | T | 6 | 99.9917 |
| 1 | 2857 | C | T | 4 | 99.99 |
| 1 | 4784 | C | T | 4 | 99.99 |
| 1 | 12885 | C | T | 4 | 99.99 |
| 1 | 17470 | C | T | 4 | 99.99 |
| 1 | 9891 | C | T | 5 | 99.988 |
| 1 | 11417 | G | T | 4 | 99.9875 |
| 1 | 27612 | C | T | 44 | 99.9866 |
| 1 | 18246 | C | T | 4 | 99.985 |
| 1 | 4936 | G | A | 47 | 99.9849 |
| 1 | 16912 | G | T | 4 | 99.9825 |
| 1 | 17321 | C | T | 4 | 99.9825 |
| 1 | 18496 | A | G | 4 | 99.9825 |
| 1 | 14408 | C | T | 178 | 99.9824 |
| 1 | 832 | C | T | 1 | 99.98 |
| 1 | 9943 | C | T | 4 | 99.98 |
| 1 | 25169 | C | T | 1 | 99.98 |
| 1 | 26367 | G | C | 2 | 99.98 |
| 1 | 23188 | C | T | 26 | 99.9788 |
| 1 | 5700 | C | A | 47 | 99.9785 |
| 1 | 27110 | C | T | 38 | 99.9784 |
| 1 | 12312 | C | T | 13 | 99.9777 |
| 1 | 18186 | G | A | 19 | 99.9758 |
| 1 | 3087 | T | C | 2 | 99.975 |
| 1 | 10039 | C | T | 2 | 99.975 |
| 1 | 23403 | A | G | 178 | 99.9749 |
| 1 | 23896 | C | T | 47 | 99.9749 |
| 1 | 17808 | G | T | 19 | 99.9747 |
| 1 | 24794 | G | A | 26 | 99.9742 |
| 1 | 8683 | C | T | 20 | 99.9735 |
| 1 | 16887 | C | T | 3 | 99.9733 |
| 1 | 18412 | G | T | 4 | 99.9725 |
| 1 | 8917 | C | T | 121 | 99.9707 |
| 1 | 18647 | C | T | 4 | 99.97 |
| 1 | 28907 | G | T | 1 | 99.97 |
| 1 | 1218 | C | T | 38 | 99.9697 |
| 1 | 3037 | C | T | 178 | 99.9693 |
| 1 | 1094 | T | C | 9 | 99.9678 |
| 1 | 27384 | T | C | 38 | 99.9661 |
| 1 | 4904 | G | T | 2 | 99.965 |
| 1 | 21071 | A | G | 2 | 99.965 |
| 1 | 25563 | G | T | 6 | 99.965 |
| 1 | 529 | G | T | 26 | 99.9604 |
| 1 | 500 | G | A | 1 | 99.96 |
| 1 | 17462 | G | A | 1 | 99.96 |

| Chr | Pos | Ref | Alt | Occurrence | Average Variation Percentage |
|---|---|---|---|---|---|
| 1 | 17479 | G | A | 1 | 99.96 |
| 1 | 20940 | G | T | 1 | 99.96 |
| 1 | 25281 | G | A | 1 | 99.96 |
| 1 | 11653 | C | T | 7 | 99.9571 |
| 1 | 25621 | G | T | 3 | 99.9567 |
| 1 | 28948 | C | T | 3 | 99.9567 |
| 1 | 241 | C | T | 177 | 99.9559 |
| 1 | 9389 | G | A | 69 | 99.9555 |
| 1 | 14857 | G | T | 2 | 99.955 |
| 1 | 17002 | C | T | 4 | 99.955 |
| 1 | 2453 | C | T | 12 | 99.9517 |
| 1 | 21077 | C | T | 2 | 99.95 |
| 1 | 18877 | C | T | 6 | 99.9483 |
| 1 | 5724 | C | T | 2 | 99.945 |
| 1 | 21486 | T | C | 4 | 99.945 |
| 1 | 1426 | C | T | 4 | 99.9425 |
| 1 | 872 | G | A | 1 | 99.94 |
| 1 | 25812 | T | C | 14 | 99.9379 |
| 1 | 186 | C | T | 8 | 99.935 |
| 1 | 20578 | G | T | 2 | 99.935 |
| 1 | 11461 | C | T | 2 | 99.93 |
| 1 | 13414 | T | C | 1 | 99.93 |
| 1 | 16323 | C | T | 2 | 99.93 |
| 1 | 27294 | C | T | 1 | 99.93 |
| 1 | 313 | C | T | 47 | 99.9289 |
| 1 | 4144 | G | A | 19 | 99.9216 |
| 1 | 20569 | G | T | 1 | 99.92 |
| 1 | 20413 | T | C | 21 | 99.9186 |
| 1 | 6355 | A | G | 1 | 99.91 |
| 1 | 9448 | C | T | 1 | 99.91 |
| 1 | 20703 | C | T | 1 | 99.91 |
| 1 | 21008 | C | T | 1 | 99.91 |
| 1 | 26933 | A | G | 1 | 99.87 |
| 1 | 26113 | G | T | 1 | 99.86 |
| 1 | 683 | C | T | 1 | 99.85 |
| 1 | 11195 | C | T | 47 | 99.8455 |
| 1 | 28899 | G | T | 2 | 99.84 |
| 1 | 5822 | C | T | 5 | 99.782 |
| 1 | 6294 | T | C | 6 | 99.7683 |
| 1 | 27092 | C | T | 1 | 99.74 |
| 1 | 16188 | G | T | 1 | 99.71 |
| 1 | 4084 | C | T | 1 | 99.69 |
| 1 | 27643 | C | A | 5 | 99.642 |
| 1 | 936 | C | T | 2 | 99.565 |
| 1 | 11619 | T | C | 2 | 99.515 |
| 1 | 6070 | C | T | 1 | 99.48 |
| 1 | 22032 | T | C | 2 | 99.435 |
| 1 | 3871 | G | T | 45 | 98.6353 |
| 1 | 25703 | C | T | 26 | 98.5604 |
| 1 | 18008 | A | G | 1 | 98.54 |
| 1 | 16750 | C | T | 8 | 98.4438 |
| 1 | 4201 | G | A | 5 | 97.268 |
| 1 | 13085 | G | A | 1 | 97.06 |
| 1 | 5907 | C | T | 1 | 96.4 |
| 1 | 3787 | C | T | 1 | 95.92 |
| 1 | 14874 | G | T | 2 | 95.845 |
| 1 | 21933 | C | T | 1 | 95.84 |
| 1 | 27493 | C | T | 1 | 95.67 |
| 1 | 21974 | G | T | 2 | 95.335 |
| 1 | 27213 | C | T | 4 | 95.29 |
| 1 | 28085 | G | T | 1 | 95.11 |
| 1 | 29675 | C | T | 4 | 94.88 |
| 1 | 11083 | G | T | 7 | 93.5314 |
| 1 | 10448 | C | T | 2 | 91.355 |
| 1 | 16557 | T | C | 1 | 88.17 |
| 1 | 19224 | T | C | 1 | 87.73 |
| 1 | 5413 | C | T | 4 | 84.415 |
| 1 | 11866 | C | T | 1 | 74.58 |
| 1 | 22675 | C | T | 1 | 63.18 |
| 1 | 19153 | A | G | 1 | 62.14 |
| 1 | 21855 | C | T | 1 | 60.22 |
| 1 | 1580 | G | T | 1 | 58.05 |
| 1 | 28086 | G | T | 1 | 50.57 |
| 1 | 15546 | C | A | 4 | 50.04 |
| **Variants with insufficient variation percentage** | | | | | |
| 1 | 14178 | C | T | 1 | 46.77 |
| 1 | 12473 | C | T | 1 | 45.72 |
| 1 | 1519 | C | T | 1 | 44.96 |
| 1 | 28895 | G | T | 1 | 44.51 |
| 1 | 337 | C | T | 46 | 44.4535 |
| 1 | 24374 | C | T | 1 | 44.05 |
| 1 | 27672 | T | C | 1 | 42.19 |
| 1 | 10465 | G | A | 1 | 42.02 |
| 1 | 5051 | C | T | 1 | 40.32 |
| 1 | 5974 | C | T | 1 | 39.5 |
| 1 | 13517 | C | T | 1 | 39.01 |
| 1 | 4148 | G | T | 1 | 35.32 |
| 1 | 1437 | C | T | 1 | 31.9 |
| 1 | 884 | C | T | 1 | 31.18 |
| 1 | 27625 | C | T | 1 | 30.26 |
| 1 | 21637 | C | T | 1 | 28.16 |
| 1 | 25579 | A | T | 1 | 27.69 |

| Chr | Pos | Ref | Alt | Occurrence | Average Variation Percentage |
|---|---|---|---|---|---|
| 1 | 29831 | C | T | 1 | 27.59 |
| 1 | 26428 | G | T | 1 | 26.97 |
| 1 | 14982 | T | C | 1 | 24.37 |
| 1 | 3994 | G | A | 1 | 24.34 |
| 1 | 2939 | C | T | 1 | 23.84 |
| 1 | 5183 | C | T | 1 | 22.78 |
| 1 | 346 | C | T | 1 | 22.6 |
| 1 | 5290 | T | C | 1 | 21.93 |
| 1 | 631 | T | C | 1 | 20.61 |
| 1 | 15194 | C | T | 1 | 20.5 |
| 1 | 18099 | T | A | 1 | 20.29 |
| 1 | 9779 | T | C | 1 | 20.26 |
