## Supplementary Table 5 for "Initial insights into the genetic epidemiology of SARS-CoV-2 isolates from Kerala suggest local spread from limited introductions"

| Chr | Start | End | Ref | Alt | Func.refGene | Gene.refGene | GeneDetail.refG | ExonicFunc.ref | AAChange.refG | GERP++_RS | PhyoP44wayCo | PhastCons44wa | SIFT_score | SIFT_pred | UNIPROT_Disul | UNIPROT_doma | UNIPROT_glyph | UNIPROT_Trans | b_CellEpitopes | cd4_Epitopes | cd4_Epitope_Sc | cd8_Epitopes | cd8_Epitope_Sc | MPDI_Potential | MPDI_Potential | ARTIC_Primers | RT_PCR-Primer | RT_PCR-Primer | Sequencing_En | Homoplasic_po | Hypermutable_ | Otherinfo |
|---|---|---|---|---|---|---|---|---|---|---|---|---|---|---|---|---|---|---|---|---|---|---|---|---|---|---|---|---|---|---|---|---|
| 1 | 186 | 186 | C | T | upstream | ORF1a | dist=80 | . | . | 1.6 | 3.21217 | 1 | . | . | . | . | . | . | . | . | . | . | . | . | . | . | . | . | . | . | . | |
| 1 | 529 | 529 | G | T | exonic | ORF1a | . | synonymous SN | ORF1a:cds-YP_ | -2.7 | 0.677165 | 0.92126 | 1 | T | . | . | . | . | . | . | . | orf1ab_APHGHV | 0.03 | . | . | . | . | . | . | . | . | |
| 1 | 872 | 872 | G | A | exonic | ORF1a | . | nonsynonymous | ORF1a:cds-YP_ | 0.741 | 2.76482 | 1 | 0.35 | T | . | . | . | . | . | . | . | . | . | . | . | . | . | . | . | . | . | |
| 1 | 936 | 936 | C | T | exonic | ORF1a | . | nonsynonymous | ORF1a:cds-YP_ | 1.65 | 3.28673 | 1 | 0.05 | D | . | . | . | . | . | . | . | orf1ab_FIDTKR | 0.05 | . | . | . | . | . | . | . | . | |
| 1 | 1580 | 1580 | G | T | exonic | ORF1a | . | nonsynonymous | ORF1a:cds-YP_ | -1.83 | 0.154331 | 0.992126 | 0.04 | D | . | . | . | . | . | . | . | . | . | . | . | nCoV-2019_6_L | . | . | . | . | . | |
| 1 | 2523 | 2523 | C | T | exonic | ORF1a | . | nonsynonymous | ORF1a:cds-YP_ | 1.65 | 3.34816 | 0.645669 | 0.01 | D | . | . | . | . | . | . | . | . | . | . | . | nCoV-2019_9_L | . | . | . | . | . | |
| 1 | 2875 | 2875 | G | T | exonic | ORF1a | . | nonsynonymous | ORF1a:cds-YP_ | -3.3 | -0.43452 | 0.96063 | 0.59 | T | . | . | . | . | . | . | . | . | . | . | . | . | . | . | . | . | . | |
| 1 | 3087 | 3087 | T | C | exonic | ORF1a | . | nonsynonymous | ORF1a:cds-YP_ | -3.03 | -0.244354 | 0 | 0.41 | T | . | . | . | . | . | . | . | orf1ab_GDCEEE | -1 | . | . | . | . | . | . | . | . | |
| 1 | 3653 | 3653 | C | T | exonic | ORF1a | . | nonsynonymous | ORF1a:cds-YP_ | 0.72 | 1.90197 | 0.992126 | 0.01 | D | . | Macro | . | . | . | . | . | . | . | . | . | . | . | . | . | . | . | |
| 1 | 3728 | 3728 | G | A | exonic | ORF1a | . | nonsynonymous | ORF1a:cds-YP_ | 1.65 | 4.256 | 1 | 0.33 | T | . | Macro | . | . | . | orf1ab polyprotei | 15 | . | . | . | . | . | . | . | . | . | . | |
| 1 | 3787 | 3787 | C | T | exonic | ORF1a | . | synonymous SN | ORF1a:cds-YP_ | -3.3 | -1.49061 | 0.015748 | 1 | T | . | Macro | . | . | . | . | . | . | . | . | . | nCoV-2019_13_ | . | . | . | . | . | |
| 1 | 3871 | 3871 | G | T | exonic | ORF1a | . | nonsynonymous | ORF1a:cds-YP_ | -3.3 | 3.28669 | 0.992126 | 0.24 | T | . | . | . | . | . | . | . | orf1ab_SEKQVE | -1 | . | . | . | . | . | . | . | . | |
| 1 | 4144 | 4144 | G | A | exonic | ORF1a | . | synonymous SN | ORF1a:cds-YP_ | -2.53 | -0.0520236 | 0.937008 | 1 | T | . | . | . | . | . | . | . | . | . | . | . | nCoV-2019_13_ | . | . | . | . | . | |
| 1 | 4201 | 4201 | G | A | exonic | ORF1a | . | nonsynonymous | ORF1a:cds-YP_ | 1.65 | 4.256 | 1 | 0.18 | T | . | . | . | . | . | . | . | . | . | . | . | . | . | . | . | . | . | |
| 1 | 4754 | 4754 | C | T | exonic | ORF1a | . | nonsynonymous | ORF1a:cds-YP_ | -3.3 | -0.539724 | 0.716535 | 0.55 | T | . | . | . | . | . | . | . | . | . | . | . | . | . | . | . | . | . | |
| 1 | 4784 | 4784 | C | T | exonic | ORF1a | . | nonsynonymous | ORF1a:cds-YP_ | -3.18 | -0.214591 | 0.0472441 | 0.06 | T | . | . | . | . | . | . | . | orf1ab_TISLAGS | -1 | . | . | . | . | . | . | . | . | |
| 1 | 4936 | 4936 | G | A | exonic | ORF1a | . | synonymous SN | ORF1a:cds-YP_ | -3.3 | 0.679528 | 0.976378 | 1 | T | . | . | . | . | . | orf1ab polyprotei | 9.2 | orf1ab_NLKTLL | 0.08 | . | . | . | . | . | . | . | . | |
| 1 | 5413 | 5413 | C | T | exonic | ORF1a | . | synonymous SN | ORF1a:cds-YP_ | -3.3 | -0.341465 | 0.984252 | 1 | T | . | Peptidase_C16 | . | . | . | . | . | . | . | . | . | . | . | . | . | . | . | |
| 1 | 5724 | 5724 | C | T | exonic | ORF1a | . | nonsynonymous | ORF1a:cds-YP_ | 0.701 | 1.5908 | 0.913386 | 0.28 | T | . | Peptidase_C16 | . | . | . | . | . | . | . | . | . | . | . | . | . | . | . | |
| 1 | 5812 | 5812 | C | G | exonic | ORF1a | . | nonsynonymous | ORF1a:cds-YP_ | -3.3 | -0.408094 | 0.984252 | 0 | D | . | Peptidase_C16 | . | . | . | . | . | . | . | . | . | . | . | . | . | . | . | |
| 1 | 5833 | 5833 | C | T | exonic | ORF1a | . | synonymous SN | ORF1a:cds-YP_ | -0.546 | 0.724614 | 0.992126 | 0.32 | T | . | Peptidase_C16 | . | . | . | . | . | orf1ab_SEYKGP | -1 | . | . | . | . | . | . | . | . | |
| 1 | 5866 | 5866 | C | T | exonic | ORF1a | . | synonymous SN | ORF1a:cds-YP_ | 1.65 | 3.32318 | 1 | 0.48 | T | . | Peptidase_C16 | . | . | . | . | . | orf1ab_SEYKGP | -1 | . | . | . | . | . | . | . | . | |
| 1 | 5907 | 5907 | C | T | exonic | ORF1a | . | nonsynonymous | ORF1a:cds-YP_ | 0.776 | 1.92395 | 0.826772 | 0.04 | D | . | Peptidase_C16 | . | . | . | . | . | orf1ab_SYTTTIK | -1 | . | . | . | . | . | . | . | . | |
| 1 | 6070 | 6070 | C | T | exonic | ORF1a | . | synonymous SN | ORF1a:cds-YP_ | -2.5 | -0.408094 | 0.023622 | 0.13 | T | . | . | . | . | . | orf1ab polyprotei | 16 | . | . | . | . | . | . | . | . | . | . | |
| 1 | 6294 | 6294 | T | C | exonic | ORF1a | . | nonsynonymous | ORF1a:cds-YP_ | 1.61 | 2.12403 | 1 | 0 | D | . | . | . | . | . | . | . | orf1ab_TYKPNT | 0.07 | . | . | . | . | . | . | . | . | |
| 1 | 6355 | 6355 | A | G | exonic | ORF1a | . | synonymous SN | ORF1a:cds-YP_ | 1.61 | 2.23063 | 1 | 0.88 | T | . | . | . | . | . | . | . | . | . | . | . | . | . | . | . | . | . | |
| 1 | 9448 | 9448 | C | T | exonic | ORF1a | . | synonymous SN | ORF1a:cds-YP_ | -3.3 | -0.224827 | 0.992134 | 0.14 | T | . | . | . | Transmembrane | . | orf1ab polyprotei | 17 | . | . | . | . | . | . | . | . | . | . | |
| 1 | 9707 | 9707 | A | G | exonic | ORF1a | . | nonsynonymous | ORF1a:cds-YP_ | 1.65 | 2.23647 | 1 | 1 | T | . | . | . | Transmembrane | . | orf1ab polyprotei | 7.3 | . | . | . | . | . | . | . | . | . | . | |
| 1 | 9891 | 9891 | C | T | exonic | ORF1a | . | nonsynonymous | ORF1a:cds-YP_ | 1.65 | 3.30935 | 1 | 0.01 | D | . | . | . | . | . | . | . | orf1ab_TQYNRY | 0.04 | . | . | . | . | . | . | . | . | |
| 1 | 9943 | 9943 | C | T | exonic | ORF1a | . | synonymous SN | ORF1a:cds-YP_ | -0.78 | 0.406276 | 1 | 1 | T | . | . | . | . | . | . | . | orf1ab_GAMDTT | 0.09 | . | . | . | . | . | . | . | . | |
| 1 | 11195 | 11195 | C | T | exonic | ORF1a | . | nonsynonymous | ORF1a:cds-YP_ | 0.68 | 1.60466 | 1 | 0.03 | D | . | . | . | Transmembrane | . | orf1ab polyprotei | 18 | orf1ab_FLLPSL | -1 | . | . | . | . | . | . | . | . | |
| 1 | 11461 | 11461 | C | T | exonic | ORF1a | . | synonymous SN | ORF1a:cds-YP_ | 0.756 | 1.8523 | 1 | 0.8 | T | . | . | . | Transmembrane | . | orf1ab polyprotei | 9.8 | orf1ab_NALDQA | 0.04 | . | . | . | . | . | . | . | . | |
| 1 | 11619 | 11619 | T | C | exonic | ORF1a | . | nonsynonymous | ORF1a:cds-YP_ | 0.411 | 0.890819 | 1 | 0.25 | T | . | . | . | Transmembrane | . | . | . | . | . | . | . | . | . | . | . | . | . | |
| 1 | 11950 | 11950 | C | T | exonic | ORF1a | . | synonymous SN | ORF1a:cds-YP_ | -0.159 | 0.684787 | 1 | 1 | T | . | . | . | . | . | . | . | . | . | . | . | . | . | . | . | . | . | |
| 1 | 12017 | 12017 | T | G | exonic | ORF1a | . | nonsynonymous | ORF1a:cds-YP_ | -3.3 | -0.208016 | 0.0745827 | 0 | D | . | . | . | . | . | orf1ab polyprotei | 18 | . | . | . | . | . | . | . | . | . | . | |
| 1 | 12325 | 12325 | C | T | exonic | ORF1a | . | synonymous SN | ORF1a:cds-YP_ | 0.787 | 1.97211 | 1 | 1 | T | . | . | . | . | . | . | . | . | . | . | . | . | . | . | . | . | . | |
| 1 | 13085 | 13085 | G | A | exonic | ORF1a | . | nonsynonymous | ORF1a:cds-YP_ | 1.65 | 4.256 | 1 | 0 | D | . | . | . | . | . | orf1ab polyprotei | 12 | orf1ab_FAVDAA | -1 | . | . | nCoV-2019_43_ | . | . | . | . | . | |
| 1 | 13414 | 13414 | T | C | exonic | ORF1a | . | synonymous SN | ORF1a:cds-YP_ | 1.65 | 2.19337 | 1 | 1 | T | . | . | . | . | . | . | . | . | . | . | . | . | . | . | . | . | . | |
| 1 | 13957 | 13957 | C | T | exonic | ORF1b | . | nonsynonymous | ORF1b:cds-YP_ | 1.65 | 3.31844 | 1 | 0.06 | T | . | . | . | . | . | orf1ab polyprotei | 15 | orf1ab_FVENPD | -1 | . | . | . | . | . | . | . | . | |
| 1 | 14120 | 14120 | C | T | exonic | ORF1b | . | nonsynonymous | ORF1b:cds-YP_ | -0.127 | 1.19331 | 1 | 0 | D | . | . | . | . | . | . | . | . | . | . | . | . | SARS-CoV-2_IB | Korea | . | . | . | |
| 1 | 14857 | 14857 | G | T | exonic | ORF1b | . | nonsynonymous | ORF1b:cds-YP_ | 0.741 | 2.74953 | 1 | 0.06 | T | . | . | . | . | . | . | . | orf1ab_FVVEVV | 0.06 | . | . | . | . | . | . | . | . | |
| 1 | 14874 | 14874 | G | T | exonic | ORF1b | . | nonsynonymous | ORF1b:cds-YP_ | -3.3 | 0.0713543 | 0.992646 | 0.01 | D | . | . | . | . | . | . | . | orf1ab_FVVEVV | 0.06 | . | . | nCoV-2019_50_ | . | . | . | . | . | |
| 1 | 15546 | 15546 | C | A | exonic | ORF1b | . | synonymous SN | ORF1b:cds-YP_ | -3.3 | -0.805717 | 0.945323 | 1 | T | . | RdRp_catalytic | . | . | . | . | . | . | . | . | . | nCoV-2019_51_ | SARS-CoV-2_IB | Korea | . | . | . | |
| 1 | 16188 | 16188 | G | T | exonic | ORF1b | . | nonsynonymous | ORF1b:cds-YP_ | 1.65 | 4.256 | 1 | 0.03 | D | . | . | . | . | . | . | . | . | . | . | . | nCoV-2019_53_ | . | . | . | . | . | |
| 1 | 16402 | 16402 | G | T | exonic | ORF1b | . | nonsynonymous | ORF1b:cds-YP_ | 1.65 | 4.256 | 1 | 0.02 | D | . | CV_ZBD | . | . | . | . | . | orf1ab_DVTDVT | -1 | . | . | . | . | . | . | . | . | |
| 1 | 16557 | 16557 | T | C | exonic | ORF1b | . | synonymous SN | ORF1b:cds-YP_ | -0.78 | 0.324472 | 1 | 1 | T | . | . | . | . | . | . | . | . | . | . | . | . | . | . | . | . | . | |
| 1 | 17462 | 17462 | G | A | exonic | ORF1b | . | nonsynonymous | ORF1b:cds-YP_ | 1.65 | 4.256 | 1 | 0 | D | . | #ERROR! | . | . | . | . | . | orf1ab_APRTLL | -1 | . | . | . | . | . | . | . | . | |
| 1 | 17470 | 17470 | C | T | exonic | ORF1b | . | synonymous SN | ORF1b:cds-YP_ | 1.65 | 3.32576 | 1 | 1 | T | . | #ERROR! | . | . | . | . | . | orf1ab_APRTLL | -1 | . | . | . | . | . | . | . | . | |
| 1 | 17479 | 17479 | G | A | exonic | ORF1b | . | nonsynonymous | ORF1b:cds-YP_ | 1.65 | 4.256 | 1 | 0 | D | . | #ERROR! | . | . | . | . | . | orf1ab_APRTLL | -1 | . | . | . | . | . | . | . | . | |
| 1 | 17808 | 17808 | G | T | exonic | ORF1b | . | nonsynonymous | ORF1b:cds-YP_ | -0.793 | 1.00017 | 1 | 0.01 | D | . | #ERROR! | . | . | . | orf1ab polyprotei | 17 | orf1ab_SPYNSQ | 0.11 | . | . | . | . | . | . | . | . | |
| 1 | 18008 | 18008 | A | G | exonic | ORF1b | . | nonsynonymous | ORF1b:cds-YP_ | 1.65 | 2.24049 | 1 | 0.23 | T | . | #ERROR! | . | . | . | . | . | . | . | . | . | . | . | . | . | . | . | |
| 1 | 18246 | 18246 | C | T | exonic | ORF1b | . | synonymous SN | ORF1b:cds-YP_ | 1.65 | 3.32576 | 1 | 1 | T | . | . | . | . | . | . | . | . | . | . | . | . | . | . | . | . | . | |
| 1 | 18496 | 18496 | A | G | exonic | ORF1b | . | nonsynonymous | ORF1b:cds-YP_ | 1.65 | 2.21939 | 1 | 0.01 | D | . | . | . | . | . | orf1ab polyprotei | 19 | . | . | . | . | . | . | . | . | . | . | |
| 1 | 18647 | 18647 | C | T | exonic | ORF1b | . | nonsynonymous | ORF1b:cds-YP_ | 1.65 | 3.27843 | 1 | 0.09 | T | . | . | . | . | . | . | . | . | . | . | . | . | . | . | . | . | . | |
| 1 | 19153 | 19153 | A | G | exonic | ORF1b | . | nonsynonymous | ORF1b:cds-YP_ | 1.65 | 2.21939 | 1 | 0.78 | T | . | . | . | . | . | orf1ab polyprotei | 17 | orf1ab_SYATHS | -1 | . | . | . | . | . | . | . | . | |
| 1 | 19224 | 19224 | T | C | exonic | ORF1b | . | synonymous SN | ORF1b:cds-YP_ | -0.794 | 0.427165 | 1 | 1 | T | . | . | . | . | . | . | . | orf1ab_YPANSI | 0.08 | . | . | nCoV-2019_64_ | . | . | . | . | . | |
| 1 | 20302 | 20302 | G | A | exonic | ORF1b | . | nonsynonymous | ORF1b:cds-YP_ | 1.65 | 4.256 | 0.984252 | 0.1 | T | . | . | . | . | . | . | . | orf1ab_RYKLEG | -1 | . | . | . | . | . | . | . | . | |
| 1 | 20413 | 20413 | T | C | exonic | ORF1b | . | synonymous SN | ORF1b:cds-YP_ | -3.3 | -0.0514961 | 0.88189 | 0.49 | T | . | . | . | . | . | orf1ab polyprotei | 20 | orf1ab_FELEDF | 0.08 | . | . | . | . | . | . | . | . | |
| 1 | 20569 | 20569 | G | T | exonic | ORF1b | . | nonsynonymous | ORF1b:cds-YP_ | 1.65 | 4.256 | 1 | 0 | D | . | . | . | . | . | orf1ab polyprotei | 18 | . | . | . | . | nCoV-2019_67_ | . | . | . | . | . | |
| 1 | 20578 | 20578 | G | T | exonic | ORF1b | . | nonsynonymous | ORF1b:cds-YP_ | 1.65 | 4.256 | 1 | 0.01 | D | . | . | . | . | . | . | . | . | . | . | . | . | . | . | . | . | . | |
| 1 | 20703 | 20703 | C | T | exonic | ORF1b | . | synonymous SN | ORF1b:cds-YP_ | -3.3 | -3.70061 | 0 | 1 | T | . | . | . | . | . | . | . | orf1ab_GVAMPN | -1 | . | . | . | . | . | . | . | . | |
| 1 | 20940 | 20940 | G | T | exonic | ORF1b | . | synonymous SN | ORF1b:cds-YP_ | 0.482 | 0.730921 | 1 | 0.38 | T | . | . | . | . | . | . | . | . | . | . | . | . | . | . | . | . | . | |
| 1 | 21008 | 21008 | C | T | exonic | ORF1b | . | nonsynonymous | ORF1b:cds-YP_ | 1.65 | 3.34955 | 0.937008 | 0 | D | . | . | . | . | . | . | . | orf1ab_TLIGDC | 0.09 | . | . | . | . | . | . | . | . | |
| 1 | 21017 | 21017 | C | T | exonic | ORF1b | . | nonsynonymous | ORF1b:cds-YP_ | 1.65 | 3.34955 | 1 | 0.93 | T | . | . | . | . | . | . | . | . | . | . | . | . | . | . | . | . | . | |
| 1 | 21071 | 21071 | A | G | exonic | ORF1b | . | nonsynonymous | ORF1b:cds-YP_ | 1.65 | 2.24167 | 1 | 0.61 | T | . | . | . | . | . | . | . | . | . | . | . | . | . | . | . | . | . | |
| 1 | 21077 | 21077 | C | T | exonic | ORF1b | . | nonsynonymous | ORF1b:cds-YP_ | -2.44 | 0.428772 | 0.440945 | 0.11 | T | . | . | . | . | . | . | . | . | . | . | . | nCoV-2019_70_ | . | . | . | . | . | |
| 1 | 21486 | 21486 | T | C | exonic | ORF1b | . | synonymous SN | ORF1b:cds-YP_ | -3.3 | -0.779827 | 0.023622 | 0.41 | T | . | . | . | . | . | orf1ab polyprotei | 11 | . | . | . | . | . | . | . | . | . | . | |
| 1 | 21621 | 21621 | C | T | exonic | S | . | nonsynonymous | S:cds-YP_00972 | -3.1 | -5.19735 | 0 | 0.12 | T | disulf_bond | BetaCoV_S1-NT | . | . | Surface glycopro | . | . | surface glycoprot | 0.12 | . | . | . | . | . | . | . | . | |
| 1 | 21846 | 21846 | C | T | exonic | S | . | nonsynonymous | S:cds-YP_00972 | -0.152 | 0.679055 | 0 | 0.07 | T | disulf_bond | BetaCoV_S1-NT | . | . | . | . | . | surface glycoprot | -1 | . | . | . | . | . | . | . | . | |
| 1 | 21933 | 21933 | C | T | exonic | S | . | nonsynonymous | S:cds-YP_00972 | -0.133 | 0.679055 | 0.00787402 | 0.41 | T | disulf_bond | BetaCoV_S1-NT | . | . | Surface glycopro | surface glycoprot | 14 | . | . | . | . | . | . | . | . | . | . | |
| 1 | 21952 | 21952 | C | T | exonic | S | . | synonymous SN | S:cds-YP_00972 | -3.3 | -2.51465 | 0 | 1 | T | disulf_bond | BetaCoV_S1-NT | . | . | . | surface glycoprot | 14 | . | . | . | . | . | . | . | . | . | . | |
| 1 | 22032 | 22032 | T | C | exonic | S | . | nonsynonymous | S:cds-YP_00972 | -0.866 | 1.0623 | 0.00787402 | 0.52 | T | disulf_bond | BetaCoV_S1-NT | . | . | Surface glycopro | surface glycoprot | 18 | . | . | . | . | nCoV-2019_72_ | . | . | . | . | . | |
| 1 | 22201 | 22201 | G | T | exonic | S | . | synonymous SN | S:cds-YP_00972 | -3.3 | -4.17537 | 0 | 0.51 | T | . | BetaCoV_S1-NT | . | . | . | . | . | surface glycoprot | 0.08 | . | . | . | . | . | . | . | . | |
| 1 | 22448 | 22448 | C | A | exonic | S | . | nonsynonymous | S:cds-YP_00972 | -0.152 | 0.679055 | 0.00787402 | 0.29 | T | disulf_bond | BetaCoV_S1-NT | . | . | . | . | . | . | . | . | . | . | . | . | . | . | . | |
| 1 | 23719 | 23719 | T | C | exonic | S | . | synonymous SN | S:cds-YP_00972 | -3.3 | -0.235276 | 0.944882 | 1 | T | . | . | . | . | Surface glycopro | . | . | surface glycoprot | 0.11 | . | . | . | . | . | . | . | . | |
| 1 | 24794 | 24794 | G | A | exonic | S | . | nonsynonymous | S:cds-YP_00972 | 1.65 | 4.256 | 0.850394 | 0.35 | T | . | . | . | . | . | . | . | . | . | . | . | . | . | . | . | . | . | |
| 1 | 25169 | 25169 | C | T | exonic | S | . | nonsynonymous | S:cds-YP_00972 | -0.194 | -0.113244 | 0.944882 | 0.01 | D | . | . | . | . | . | . | . | surface glycoprot | -1 | Spike glycoprote | KNHTSPDVDLG | . | . | . | . | . | . | |
| 1 | 25281 | 25281 | G | A | exonic | S | . | nonsynonymous | S:cds-YP_00972 | 1.65 | 4.256 | 1 | 0 | D | . | . | . | . | . | . | . | . | . | . | . | nCoV-2019_84_ | . | . | . | . | . | |
| 1 | 25546 | 25546 | C | T | exonic | ORF3a | . | nonsynonymous | ORF3a:cds-YP_ | -3.3 | -1.39205 | 0.685039 | 0.27 | T | . | . | . | Transmembrane | . | ORF3a protein_A | 11 | . | . | . | . | . | . | . | . | . | . | |
| 1 | 25703 | 25703 | C | T | exonic | ORF3a | . | nonsynonymous | ORF3a:cds-YP_ | 0.597 | 0.612417 | 0.732283 | 0.23 | T | . | . | . | Transmembrane | . | . | . | ORF3a protein_ | -1 | . | . | . | . | . | . | . | . | |
| 1 | 25812 | 25812 | T | C | exonic | ORF3a | . | synonymous SN | ORF3a:cds-YP_ | -3.3 | -3.77857 | 0.023622 | 1 | T | . | . | . | . | . | ORF3a protein_K | 13 | ORF3a protein_N | 0.03 | . | . | . | . | . | . | . | . | |
| 1 | 26113 | 26113 | G | T | exonic | ORF3a | . | stopgain | ORF3a:cds-YP_ | 0.154 | 0.238717 | 0 | . | . | . | . | . | . | . | . | . | ORF3a protein_E | -1 | . | . | . | . | . | . | . | . | |
| 1 | 26367 | 26367 | G | C | exonic | E | . | synonymous SN | E:cds-YP_00972 | 0.715 | 2.57435 | 1 | 1 | T | . | Intravirion | . | . | . | envelope protein | 15 | . | . | . | . | . | SARS-CoV-2_IB | Korea | . | . | . | |
| 1 | 26486 | 26486 | T | A | upstream;downst | M;ORF6;ORF7a; | dist=37;dist=14 | . | . | 1.65 | 2.20065 | 1 | . | . | . | . | . | . | . | . | . | . | . | . | . | . | . | . | . | . | . | |
| 1 | 26933 | 26933 | A | G | exonic | M | . | synonymous SN | M:cds-YP_0097 | 1.65 | 2.20759 | 1 | 1 | T | . | Intravirion | . | . | . | membrane glyco | 20 | membrane glyco | 0.08 | . | . | . | . | . | . | . | . | |
| 1 | 27643 | 27643 | C | A | exonic | ORF7a | . | nonsynonymous | ORF7a:cds-YP_ | 1.65 | 3.2857 | 1 | 0 | D | . | . | . | . | . | ORF7a protein_V | 15 | ORF7a protein_F | 0.1 | . | . | . | . | . | . | . | . | |
| 1 | 28024 | 28024 | A | G | exonic | ORF8 | . | nonsynonymous | ORF8:cds-YP_0 | -0.11 | 0.731118 | 0 | 0.16 | T | . | . | . | . | . | ORF8 protein_FY | 13 | . | . | . | . | . | . | . | . | . | . | |
| 1 | 28028 | 28028 | G | A | exonic | ORF8 | . | stopgain | ORF8:cds-YP_0 | 0.372 | 1.22069 | 0.165354 | . | . | . | . | . | . | . | ORF8 protein_FY | 13 | . | . | . | . | . | . | . | . | . | . | |
| 1 | 29659 | 29659 | C | T | exonic | ORF10 | . | synonymous SN | ORF10:cds-YP_ | 1.6 | 3.25353 | 1 | . | . | . | . | . | . | . | . | . | . | . | . | . | . | . | . | . | . | . | |
