## Supplementary Table 6 for "Initial insights into the genetic epidemiology of SARS-CoV-2 isolates from Kerala suggest local spread from limited introductions"

Number of variable sites: 592

=========== Haplotype Distribution ===========

Number of haplotypes, h: 400

Haplotype diversity, Hd: 0.9651

| | |
|---|---|
| Hap_1: 1 | [India/NIV-38793/2020] |
| Hap_2: 144 | [India/NIV-40837/2020 India/NIV-10176/2020 India/NIV-14614/2020 India/NIV-13307/2020 India/MW26/2020 India/GA38/2020 India/GA67/2020 India/GA83/2020 India/GA46/2020 India/GA50/2020 India/GA51/2020 India/GA47/2020 India/GA49/2020 India/GA57/2020 India/GA60/2020 India/GA98/2020 India/GA8/2020 India/GA41/2020 India/GA59/2020 India/GA74/2020 India/GA39/2020 India/GA68/2020 India/TF38/2020 India/GBRC56 |
| Hap_3: 1 | [India/NIV-12628/2020] |
| Hap_4: 2 | [India/NIV-11670/2020 India/NIV-12067/2020] |
| Hap_5: 1 | [India/GA78/2020] |
| Hap_6: 1 | [India/GA79/2020] |
| Hap_7: 1 | [India/TF3/2020] |
| Hap_8: 3 | [India/GA61/2020 India/GA80/2020 India/GA63/2020] |
| Hap_9: 2 | [India/GA54/2020 India/GA55/2020] |
| Hap_10: 6 | [India/GA45/2020 India/GA62/2020 India/GA82/2020 India/GA81/2020 India/GA9/2020 India/GA30/2020] |
| Hap_11: 1 | [India/GA5/2020] |
| Hap_12: 1 | [India/GA53/2020] |
| Hap_13: 1 | [India/GA52/2020] |
| Hap_14: 1 | [India/GA56/2020] |
| Hap_15: 1 | [India/GA42/2020] |
| Hap_16: 1 | [India/GA48/2020] |
| Hap_17: 1 | [India/GA87/2020] |
| Hap_18: 1 | [India/GA88/2020] |
| Hap_19: 1 | [India/GA40/2020] |
| Hap_20: 1 | [India/GA6/2020] |
| Hap_21: 1 | [India/GA85/2020] |
| Hap_22: 2 | [India/MW33/2020 India/MW32/2020] |
| Hap_23: 1 | [India/GA43/2020] |
| Hap_24: 1 | [India/GA37/2020] |
| Hap_25: 1 | [India/GA77/2020] |
| Hap_26: 2 | [India/GA75/2020 India/GA76/2020] |
| Hap_27: 2 | [India/GA69/2020 India/GA70/2020] |
| Hap_28: 1 | [India/GA7/2020] |
| Hap_29: 2 | [India/MW28/2020 India/MW29/2020] |
| Hap_30: 1 | [India/GA36/2020] |
| Hap_31: 31 | [India/IB6/2020 India/IB11/2020 India/InStem_NCBS_0082/2020 India/InStem_NCBS_0074/2020 India/CCMB_C9/2020 India/CCMB_C16/2020 India/InStem_NCBS_0040/2020 India/InStem_NCBS_0037/2020 India/InStem_NCBS_0014/2020 India/InStem_NCBS_0028/2020 India/IB12/2020 India/IB13/2020 India/IB15/2020 India/IB21/2020 India/IB16/2020 India/IB18/2020 India/IB2/2020 India/IB17/2020 India/IB26/2020 India/IB31/2020 |
| Hap_32: 1 | [India/IB62/2020] |
| Hap_33: 1 | [India/IB7/2020] |
| Hap_34: 1 | [India/IB1/2020] |
| Hap_35: 1 | [India/IB9/2020] |
| Hap_36: 1 | [India/IB8/2020] |
| Hap_37: 1 | [India/GA71/2020] |
| Hap_38: 1 | [India/IB10/2020] |
| Hap_39: 1 | [India/GA86/2020] |
| Hap_40: 1 | [India/GA66/2020] |
| Hap_41: 1 | [India/GA4/2020] |
| Hap_42: 1 | [India/GBRC251a/2020] |
| Hap_43: 1 | [India/GBRC274/2020] |
| Hap_44: 1 | [India/TF26/2020] |
| Hap_45: 1 | [India/TF4/2020] |
| Hap_46: 1 | [India/TF6/2020] |
| Hap_47: 2 | [India/MW34/2020 India/MW36/2020] |
| Hap_48: 1 | [India/MN1/2020] |
| Hap_49: 3 | [India/TF41/2020 India/NCDC6082_CSIR-IGIB/2020 India/NCDC4462_CSIR-IGIB/2020] |
| Hap_50: 1 | [India/TF30/2020] |
| Hap_51: 1 | [India/TF7/2020] |
| Hap_52: 1 | [India/NIV-10419/2020] |
| Hap_53: 1 | [India/NIV-11687/2020] |
| Hap_54: 1 | [India/NIV-11855/2020] |
| Hap_55: 1 | [India/NIV-12019/2020] |
| Hap_56: 1 | [India/NIV-11677/2020] |
| Hap_57: 4 | [India/CCMB_L1011/2020 India/CCMB_L1023/2020 India/CCMB_L1021/2020 India/CCMB_L1024/2020] |
| Hap_58: 37 | [India/CCMB_L1824_ONT/2020 India/CCMB_L1885/2020 India/CCMB_OM1/2020 India/CCMB_OM11/2020 India/CCMB_OM12/2020 India/CCMB_OM13/2020 India/CCMB_OM16/2020 India/CCMB_OM15/2020 India/CCMB_OM19/2020 India/CCMB_OM7/2020 India/CCMB_OM17/2020 India/CCMB_OM1_ONT/2020 India/CCMB_M471/2020 India/CCMB_M686/2020 India/CCMB_M806/2020 India/CCMB_M860/2020 India/CCMB_L16 |
| Hap_59: 1 | [India/CCMB_L1934/2020] |
| Hap_60: 1 | [India/CCMB_L1822_ONT/2020] |
| Hap_61: 1 | [India/CCMB_L1893/2020] |
| Hap_62: 3 | [India/CCMB_L1864/2020 India/CCMB_M891/2020 India/CCMB_M456/2020] |
| Hap_63: 1 | [India/CCMB_M175/2020] |
| Hap_64: 1 | [India/CCMB_M2/2020] |
| Hap_65: 1 | [India/CCMB_M288/2020] |
| Hap_66: 1 | [India/CCMB_M152/2020] |
| Hap_67: 3 | [India/CCMB_M135/2020 India/CCMB_L1746/2020 India/CCMB_L1811/2020] |
| Hap_68: 1 | [India/CCMB_M22/2020] |
| Hap_69: 1 | [India/CCMB_L1892/2020] |
| Hap_70: 1 | [India/CCMB_M25/2020] |
| Hap_71: 8 | [India/CCMB_L1959/2020 India/CCMB_M170/2020 India/CCMB_L1930/2020 India/CCMB_M489/2020 India/CCMB_M717/2020 India/CCMB_M882/2020 India/CCMB_M697/2020 India/CCMB_L1694/2020] |
| Hap_72: 2 | [India/CCMB_M357/2020 India/CCMB_M441/2020] |
| Hap_73: 1 | [India/CCMB_M116/2020] |
| Hap_74: 6 | [India/CCMB_L1967/2020 India/CCMB_M785/2020 India/CCMB_L1819/2020 India/CCMB_M651/2020 India/CCMB_M466/2020 India/CCMB_M652/2020] |
| Hap_75: 1 | [India/CCMB_M14/2020] |
| Hap_76: 1 | [India/CCMB_L1931/2020] |
| Hap_77: 1 | [India/CCMB_M118/2020] |
| Hap_78: 1 | [India/CCMB_M343/2020] |
| Hap_79: 1 | [India/CCMB_M200/2020] |
| Hap_80: 1 | [India/CCMB_M151/2020] |
| Hap_81: 1 | [India/CCMB_M172/2020] |
| Hap_82: 2 | [India/CCMB_L1657/2020 India/CCMB_M465/2020] |
| Hap_83: 3 | [India/CCMB_L1963/2020 India/UN2/2020 India/CCMB_L1564/2020] |
| Hap_84: 1 | [India/CCMB_L1935/2020] |
| Hap_85: 1 | [India/DRDE_3983/2020] |

| | |
|---|---|
| Hap_86: 1 | [India/NCDC7624_CSIR-IGIB/2020] |
| Hap_87: 1 | [India/NCDC-02155/2020] |
| Hap_88: 1 | [India/GA93/2020] |
| Hap_89: 1 | [India/GA65/2020] |
| Hap_90: 1 | [India/NIV-26133/2020] |
| Hap_91: 1 | [India/IB54/2020] |
| Hap_92: 1 | [India/NIV-42694/2020] |
| Hap_93: 3 | [India/GA84/2020 India/GA11/2020 India/GA33/2020] |
| Hap_94: 1 | [India/InStem_NCBS_0062/2020] |
| Hap_95: 3 | [India/InStem_NCBS_0073/2020 India/InStem_NCBS_0076/2020 India/InStem_NCBS_0039/2020] |
| Hap_96: 1 | [India/ILSCV34939/2020] |
| Hap_97: 17 | [India/ILSCV35353/2020 India/ILSCV32898/2020 India/ILSCV35350/2020 India/ILSCV35317/2020 India/ILSCV35366/2020 India/ILSCV35333/2020 India/ILSCV35376/2020 India/ILSCV32899/2020 India/ILSCV31185/2020 India/CCMB_C6/2020 India/CCMB_C5/2020 India/CCMB_C11/2020 India/CCMB_C19/2020 India/ILSCV16809/2020 India/IB22/2020 India/IB30/2020 India/IB27/2020] |
| Hap_98: 1 | [India/ILSCV28594/2020] |
| Hap_99: 2 | [India/ILSCV32872/2020 India/ILSCV32870/2020] |
| Hap_100: 9 | [India/ILSCV32892/2020 India/ILSCV32896/2020 India/ILSCV33982/2020 India/ILSCV32895/2020 India/ILSCV32889/2020 India/ILSCV32893/2020 India/ILSCV34480/2020 India/ILSCV34478/2020 India/ILSCV33971/2020] |
| Hap_101: 1 | [India/ILSCV30442/2020] |
| Hap_102: 2 | [India/ILSCV32485/2020 India/ILSCV32486/2020] |
| Hap_103: 1 | [India/ILSCV35461/2020] |
| Hap_104: 2 | [India/ILSCV27713/2020 India/GBRC318b/2020] |
| Hap_105: 1 | [India/ILSCV35475/2020] |
| Hap_106: 1 | [India/ILSCV35351/2020] |
| Hap_107: 1 | [India/ILSCV33972/2020] |
| Hap_108: 2 | [India/ILSCV25524/2020 India/NCCS_NR2122/2020] |
| Hap_109: 1 | [India/ILSCV29997/2020] |
| Hap_110: 1 | [India/ILSCV30167/2020] |
| Hap_111: 1 | [India/ILSCV29457/2020] |
| Hap_112: 1 | [India/ILSCV35141/2020] |
| Hap_113: 1 | [India/ILSCV30115/2020] |
| Hap_114: 1 | [India/ILSCV34933/2020] |
| Hap_115: 3 | [India/ILSCV32593/2020 CS1898 India/NCCS_NR1445/2020] |
| Hap_116: 1 | [India/ILSCV25367/2020] |
| Hap_117: 5 | [India/ILSCV25591/2020 India/ILSCV25594/2020 India/CCMB_C7/2020 India/CCMB_C17/2020 India/ILSCV20557/2020] |
| Hap_118: 4 | [India/AR10/2020 India/AR12/2020 India/AR4/2020 India/AR9/2020] |
| Hap_119: 4 | [India/AR60/2020 India/AR62/2020 India/MaxCov0022_CSIR-IGIB/2020 India/MaxCov0010_CSIR-IGIB/2020] |
| Hap_120: 2 | [India/AR5/2020 India/AR8/2020] |
| Hap_121: 1 | [India/AR15/2020] |
| Hap_122: 2 | [India/AR30/2020 India/AR31/2020] |
| Hap_123: 1 | [India/AR54/2020] |
| Hap_124: 1 | [India/AR16/2020] |
| Hap_125: 2 | [India/AR25/2020 India/AR26/2020] |
| Hap_126: 1 | [India/ILSCV25518/2020] |
| Hap_127: 4 | [India/AR41/2020 India/AR40/2020 India/AR42/2020 India/AR43/2020] |
| Hap_128: 1 | [India/ILSCV27406/2020] |
| Hap_129: 5 | [India/AFMC_5306/2020 India/AFMC_5304/2020 India/AFMC_5286/2020 India/AFMC_5309/2020 India/AFMC_5318/2020] |
| Hap_130: 1 | [India/GA12/2020] |
| Hap_131: 1 | [India/GA72/2020] |
| Hap_132: 4 | [India/ILSCV32814/2020 India/ILSCV32815/2020 India/ILSCV32816/2020 India/ILSCV32817/2020] |
| Hap_133: 1 | [India/GJ-GBRC-395a/2020] |
| Hap_134: 1 | [India/GJ-GBRC-395b/2020] |
| Hap_135: 1 | [India/AR23/2020] |
| Hap_136: 2 | [India/GA18/2020 India/GA90/2020] |
| Hap_137: 1 | [India/AR27/2020] |
| Hap_138: 1 | [India/MW39/2020] |
| Hap_139: 1 | [India/AR65/2020] |
| Hap_140: 1 | [India/ILSCV31650/2020] |
| Hap_141: 2 | [India/GJ-GBRC-376a/2020 India/GJ-GBRC-376b/2020] |
| Hap_142: 2 | [CS1897 CS1795] |
| Hap_143: 7 | [CS1082 CS1090 CS1092 CS1887 CS1091 CS1801 CS1093] |
| Hap_144: 8 | [India/ILSCV35308/2020 India/CCMB_C12/2020 India/CCMB_C8/2020 India/CCMB_C10/2020 India/IB24/2020 India/IB44/2020 India/IB29/2020 India/S5/2020] |
| Hap_145: 1 | [India/AR61/2020] |
| Hap_146: 1 | [India/AR57/2020] |
| Hap_147: 2 | [India/AR13/2020 India/AR14/2020] |
| Hap_148: 1 | [India/AR3/2020] |
| Hap_149: 1 | [India/AR6/2020] |
| Hap_150: 1 | [India/AR28/2020] |
| Hap_151: 1 | [India/AR29/2020] |
| Hap_152: 2 | [India/AR24/2020 India/AR32/2020] |
| Hap_153: 1 | [India/TF21/2020] |
| Hap_154: 1 | [India/UP-NBRI-N48-TEI094/2020] |
| Hap_155: 1 | [India/UP-NBRI-N58-TEG811/2020] |
| Hap_156: 1 | [India/UP-NBRI-N55-TEH861/2020] |
| Hap_157: 1 | [India/AR59/2020] |
| Hap_158: 1 | [India/AR20/2020] |
| Hap_159: 1 | [India/AR53/2020] |
| Hap_160: 3 | [CS1903 CS1076 CS1094] |
| Hap_161: 4 | [India/KA-nimh-19696/2020 India/CCMB_M79/2020 India/CCMB_M81/2020 India/CCMB_M80/2020] |
| Hap_162: 34 | [CS1899 CS1902 CS1086 CS1901 CS1806 CS1804 CS1807 CS1872 CS1878 CS1121 CS1811 CS1137 CS1815 CS1814 CS1816 CS1817 CS1869 CS1829 CS1868 CS1819 CS1843 CS1122 CS1812 CS1821 CS1865 CS1123 CS1877 CS1846 CS1873 CS1870 CS1853 CS1855 CS1871 CS1874] |
| Hap_163: 2 | [CS1805 CS1803] |
| Hap_164: 1 | [CS1077] |
| Hap_165: 1 | [India/IB64/2020] |
| Hap_166: 2 | [India/MW22/2020 India/MW20/2020] |
| Hap_167: 1 | [India/AR44/2020] |
| Hap_168: 1 | [India/IB59/2020] |
| Hap_169: 1 | [India/MW40/2020] |
| Hap_170: 1 | [India/AR48/2020] |
| Hap_171: 1 | [India/GA95/2020] |
| Hap_172: 1 | [India/AR50/2020] |
| Hap_173: 1 | [India/GBRC327a/2020] |
| Hap_174: 1 | [India/GBRC327b/2020] |
| Hap_175: 1 | [India/AR38/2020] |
| Hap_176: 1 | [India/GA94/2020] |

| | |
|---|---|
| Hap_177: 1 | [India/InStem_NCBS_0075/2020] |
| Hap_178: 1 | [India/NIV-7832/2020] |
| Hap_179: 1 | [India/NIV-QA-708/2020] |
| Hap_180: 1 | [India/GBRC206/2020] |
| Hap_181: 1 | [India/GBRC220/2020] |
| Hap_182: 2 | [India/S75/2020 India/S74/2020] |
| Hap_183: 1 | [India/NIV-QC-796/2020] |
| Hap_184: 1 | [India/GBRC203a/2020] |
| Hap_185: 1 | [India/GBRC237a/2020] |
| Hap_186: 1 | [India/CCMB_C14/2020] |
| Hap_187: 1 | [India/CCMB_C15/2020] |
| Hap_188: 1 | [India/CCMB_C18/2020] |
| Hap_189: 1 | [India/CCMB_L1082/2020] |
| Hap_190: 6 | [India/CCMB_L1085/2020 India/ILSCV16514/2020 India/CCMB_OM10/2020 India/CCMB_M750/2020 India/CCMB_M950/2020 India/CCMB_L1554/2020] |
| Hap_191: 8 | [India/CCMB_L1100/2020 India/CCMB_L1103/2020 India/CCMB_L1108/2020 India/CCMB_OM2/2020 India/CCMB_OM5/2020 India/CCMB_OM2_ONT/2020 India/CCMB_OM6_ONT/2020 India/NGC-CDFD-01/2020] |
| Hap_192: 1 | [India/CCMB_L1089/2020] |
| Hap_193: 1 | [India/CCMB_L1090/2020] |
| Hap_194: 1 | [India/CCMB_L1017/2020] |
| Hap_195: 1 | [India/CCMB_L1026/2020] |
| Hap_196: 1 | [India/CCMB_L1022/2020] |
| Hap_197: 1 | [India/CCMB_L1025/2020] |
| Hap_198: 1 | [India/ILSCV35574/2020] |
| Hap_199: 1 | [India/ILSCV24918/2020] |
| Hap_200: 1 | [India/GBRC318a/2020] |
| Hap_201: 1 | [India/GBRC320/2020] |
| Hap_202: 1 | [India/GBRC265/2020] |
| Hap_203: 1 | [India/GBRC276/2020] |
| Hap_204: 1 | [India/GBRC281/2020] |
| Hap_205: 1 | [India/ILSCV15546/2020] |
| Hap_206: 1 | [India/ILSCV15552/2020] |
| Hap_207: 1 | [India/ILSCV20408/2020] |
| Hap_208: 1 | [India/ILSCV20407/2020] |
| Hap_209: 1 | [India/ILSCV15547/2020] |
| Hap_210: 1 | [India/InStem_NCBS_0041/2020] |
| Hap_211: 1 | [India/NIV-QA-707/2020] |
| Hap_212: 1 | [India/GBRC270/2020] |
| Hap_213: 2 | [India/NIV-17002/2020 India/NIV-23721/2020] |
| Hap_214: 1 | [India/NIV-21455/2020] |
| Hap_215: 2 | [India/NIV-15879/2020 India/NIV-12024/2020] |
| Hap_216: 1 | [India/NIV-29909/2020] |
| Hap_217: 1 | [India/NIV-28416/2020] |
| Hap_218: 3 | [India/NIV-20134/2020 India/BJMC_965/2020 India/BJMC_1123/2020] |
| Hap_219: 1 | [India/NIV-23098/2020] |
| Hap_220: 1 | [India/NIV-18650/2020] |
| Hap_221: 3 | [India/NIV-36461/2020 India/NIV-65861/2020 India/NIV-65829/2020] |
| Hap_222: 1 | [India/NIV-26115/2020] |
| Hap_223: 1 | [India/NIV-37985/2020] |
| Hap_224: 1 | [India/NIV-16503/2020] |
| Hap_225: 1 | [India/NIV-37681/2020] |
| Hap_226: 1 | [India/NIV-25950/2020] |
| Hap_227: 1 | [India/NIV-28900/2020] |
| Hap_228: 2 | [India/NIV-15351/2020 India/BJMC_2035/2020] |
| Hap_229: 1 | [India/S79/2020] |
| Hap_230: 1 | [India/S73/2020] |
| Hap_231: 1 | [India/NIV-11676/2020] |
| Hap_232: 1 | [India/NIV-12058/2020] |
| Hap_233: 1 | [India/InStem_NCBS_0032/2020] |
| Hap_234: 1 | [India/TF10/2020] |
| Hap_235: 1 | [India/S81/2020] |
| Hap_236: 1 | [India/TF15/2020] |
| Hap_237: 1 | [India/TF14/2020] |
| Hap_238: 1 | [India/TF25/2020] |
| Hap_239: 1 | [India/TF2/2020] |
| Hap_240: 1 | [India/TF12/2020] |
| Hap_241: 1 | [India/TF13/2020] |
| Hap_242: 1 | [India/TF17/2020] |
| Hap_243: 2 | [India/NCDC1895_CSIR-IGIB/2020 India/BJMC_1760/2020] |
| Hap_244: 1 | [India/NCDC3832_CSIR-IGIB/2020] |
| Hap_245: 1 | [India/NCDC4402_CSIR-IGIB/2020] |
| Hap_246: 1 | [India/ILSCV21602/2020] |
| Hap_247: 1 | [India/ILSCV21721/2020] |
| Hap_248: 1 | [India/GBRC290b/2020] |
| Hap_249: 1 | [India/ILSCV19209/2020] |
| Hap_250: 1 | [India/MaxCov0012_CSIR-IGIB/2020] |
| Hap_251: 1 | [India/InStem_NCBS_0038/2020] |
| Hap_252: 1 | [India/NIV-11182/2020] |
| Hap_253: 1 | [India/InStem_NCBS_0031/2020] |
| Hap_254: 1 | [India/InStem_NCBS_0042/2020] |
| Hap_255: 1 | [India/InStem_NCBS_0035/2020] |
| Hap_256: 1 | [India/InStem_NCBS_0026/2020] |
| Hap_257: 1 | [India/InStem_NCBS_0022/2020] |
| Hap_258: 1 | [India/InStem_NCBS_0013/2020] |
| Hap_259: 1 | [India/IB19/2020] |
| Hap_260: 2 | [India/MW17/2020 India/MW18/2020] |
| Hap_261: 1 | [India/GA22/2020] |
| Hap_262: 1 | [India/GA34/2020] |
| Hap_263: 1 | [India/GA21/2020] |
| Hap_264: 1 | [India/GA19/2020] |
| Hap_265: 1 | [India/NCCS_NR2513/2020] |
| Hap_266: 1 | [India/GA16/2020] |
| Hap_267: 4 | [India/GA24/2020 India/GA25/2020 India/GA26/2020 India/GA27/2020] |

| | |
|---|---|
| Hap_268: 1 | [India/NCCS_NR1922/2020] |
| Hap_269: 2 | [India/NCCS_NR2329/2020 India/NCCS_NR2328/2020] |
| Hap_270: 1 | [India/BJMC_2030/2020] |
| Hap_271: 1 | [India/GA29/2020] |
| Hap_272: 1 | [India/GA17/2020] |
| Hap_273: 2 | [India/MN10/2020 India/MN11/2020] |
| Hap_274: 8 | [India/CCMB_OM8/2020 India/CCMB_M588/2020 India/CCMB_M900/2020 India/CCMB_M925/2020 India/CCMB_M931/2020 India/CCMB_M969/2020 India/CCMB_M971/2020 India/CCMB_M581/2020] |
| Hap_275: 3 | [India/CCMB_OM4/2020 India/CCMB_OM9/2020 India/CCMB_OM4_ONT/2020] |
| Hap_276: 1 | [India/CCMB_OM14/2020] |
| Hap_277: 1 | [India/CCMB_OM3/2020] |
| Hap_278: 1 | [India/CCMB_OM18/2020] |
| Hap_279: 1 | [India/CCMB_OM6/2020] |
| Hap_280: 1 | [India/CCMB_OM20/2020] |
| Hap_281: 2 | [India/NCCS_NR2107/2020 India/NCCS_NR2108/2020] |
| Hap_282: 1 | [India/CCMB_M739/2020] |
| Hap_283: 6 | [India/CCMB_M762/2020 India/CCMB_M780/2020 India/CCMB_M830/2020 India/CCMB_M838/2020 India/CCMB_M841/2020 India/CCMB_M772/2020] |
| Hap_284: 1 | [India/CCMB_M844/2020] |
| Hap_285: 2 | [India/CCMB_M840/2020 India/CCMB_M5/2020] |
| Hap_286: 1 | [India/CCMB_M82/2020] |
| Hap_287: 1 | [India/CCMB_M722/2020] |
| Hap_288: 1 | [India/CCMB_M842/2020] |
| Hap_289: 1 | [India/CCMB_M733/2020] |
| Hap_290: 1 | [India/CCMB_N4/2020] |
| Hap_291: 1 | [India/CCMB_M995/2020] |
| Hap_292: 1 | [India/CCMB_M696/2020] |
| Hap_293: 2 | [India/CCMB_M700/2020 India/CCMB_M711/2020] |
| Hap_294: 1 | [India/CCMB_M818/2020] |
| Hap_295: 1 | [India/CCMB_M712/2020] |
| Hap_296: 1 | [India/CCMB_M740/2020] |
| Hap_297: 1 | [India/CCMB_M892/2020] |
| Hap_298: 1 | [India/CCMB_M845/2020] |
| Hap_299: 1 | [India/CCMB_N18/2020] |
| Hap_300: 1 | [India/CCMB_M804/2020] |
| Hap_301: 1 | [India/BJMC_1769/2020] |
| Hap_302: 1 | [India/CCMB_M64/2020] |
| Hap_303: 1 | [India/nimh-0834/2020] |
| Hap_304: 1 | [India/MN32/2020] |
| Hap_305: 1 | [India/MN4/2020] |
| Hap_306: 1 | [India/MN8/2020] |
| Hap_307: 1 | [India/NCCS_NR1333/2020] |
| Hap_308: 1 | [India/nimh-0182/2020] |
| Hap_309: 1 | [India/BJMC_2086/2020] |
| Hap_310: 1 | [India/MN5/2020] |
| Hap_311: 1 | [India/NCCS_NR1474/2020] |
| Hap_312: 1 | [India/nimh-0116/2020] |
| Hap_313: 1 | [India/CCMB_C4-15/2020] |
| Hap_314: 1 | [India/IB36/2020] |
| Hap_315: 2 | [India/IB28/2020 India/IB33/2020] |
| Hap_316: 1 | [India/IB25/2020] |
| Hap_317: 1 | [India/BJMC_2192/2020] |
| Hap_318: 1 | [India/BJMC_6185/2020] |
| Hap_319: 1 | [India/IB41/2020] |
| Hap_320: 1 | [India/IB50/2020] |
| Hap_321: 1 | [India/IB48/2020] |
| Hap_322: 1 | [India/CCMB_L1809/2020] |
| Hap_323: 1 | [India/CCMB_L1724/2020] |
| Hap_324: 1 | [India/CCMB_L1726/2020] |
| Hap_325: 1 | [India/CCMB_L1728/2020] |
| Hap_326: 1 | [India/CCMB_L1675/2020] |
| Hap_327: 1 | [India/CCMB_L1810/2020] |
| Hap_328: 1 | [India/CCMB_L1738/2020] |
| Hap_329: 1 | [India/CCMB_L1781/2020] |
| Hap_330: 1 | [India/CCMB_L1806/2020] |
| Hap_331: 1 | [India/CCMB_L1743/2020] |
| Hap_332: 1 | [India/BJMC_2442/2020] |
| Hap_333: 2 | [India/NGC-CDFD-05/2020 India/NGC-CDFD-07/2020] |
| Hap_334: 1 | [India/NGC-CDFD-11/2020] |
| Hap_335: 1 | [India/NGC-CDFD-12/2020] |
| Hap_336: 1 | [India/NGC-CDFD-30/2020] |
| Hap_337: 1 | [India/NGC-CDFD-33/2020] |
| Hap_338: 2 | [India/NGC-CDFD-24/2020 India/NGC-CDFD-34/2020] |
| Hap_339: 2 | [India/S3/2020 India/S2/2020] |
| Hap_340: 1 | [India/CCMB_M66/2020] |
| Hap_341: 1 | [India/CCMB_M361/2020] |
| Hap_342: 1 | [India/CCMB_M458/2020] |
| Hap_343: 1 | [India/CCMB_M437/2020] |
| Hap_344: 1 | [India/CCMB_M446/2020] |
| Hap_345: 1 | [India/CCMB_M451/2020] |
| Hap_346: 1 | [India/CCMB_M363/2020] |
| Hap_347: 1 | [India/IB38/2020] |
| Hap_348: 1 | [India/RMRC2/2020] |
| Hap_349: 1 | [India/NIV-64478/2020] |
| Hap_350: 1 | [India/NIV-64904/2020] |
| Hap_351: 1 | [India/NIV-65282/2020] |
| Hap_352: 1 | [India/NIV-65417/2020] |
| Hap_353: 1 | [India/NIV-64905/2020] |
| Hap_354: 1 | [India/NIV-47470/2020] |
| Hap_355: 1 | [India/NIV-64850/2020] |
| Hap_356: 1 | [India/NIV-65813/2020] |
| Hap_357: 1 | [India/NIV-65835/2020] |
| Hap_358: 1 | [India/NIV-65877/2020] |

| | |
|---|---|
| Hap_359: 1 | [India/NIV-65847/2020] |
| Hap_360: 1 | [India/NIV-64908/2020] |
| Hap_361: 1 | [India/NIV-46317/2020] |
| Hap_362: 1 | [India/NIV-7830/2020] |
| Hap_363: 1 | [India/NIV-QC-802/2020] |
| Hap_364: 1 | [India/NIV-45936/2020] |
| Hap_365: 1 | [India/NIV-64877/2020] |
| Hap_366: 1 | [India/NIV-64463/2020] |
| Hap_367: 1 | [India/BJMC_1658/2020] |
| Hap_368: 2 | [India/GBRC97a/2020 India/GBRC97b/2020] |
| Hap_369: 1 | [India/NIV-46348/2020] |
| Hap_370: 1 | [India/NGC-CDFD-06/2020] |
| Hap_371: 1 | [India/AFMC_5868/2020] |
| Hap_372: 1 | [India/NGC-CDFD-04/2020] |
| Hap_373: 1 | [India/NGC-CDFD-18/2020] |
| Hap_374: 1 | [India/NGC-CDFD-21/2020] |
| Hap_375: 1 | [India/InStem_NCBS_0067/2020] |
| Hap_376: 1 | [India/AFMC_5827/2020] |
| Hap_377: 1 | [India/InStem_NCBS_0069/2020] |
| Hap_378: 11 | [CS1836 CS1837 CS1831 CS1839 CS1840 CS1847 CS1867 CS1799 CS1885 CS1798 CS1838] |
| Hap_379: 1 | [CS1800] |
| Hap_380: 2 | [CS1143 CS1144] |
| Hap_381: 1 | [CS1875] |
| Hap_382: 1 | [CS1830] |
| Hap_383: 5 | [CS1110 CS1113 CS1880 CS1848 CS1112] |
| Hap_384: 2 | [CS1824 CS1823] |
| Hap_385: 2 | [CS1159 CS1160] |
| Hap_386: 2 | [CS1842 CS1841] |
| Hap_387: 7 | [CS1129 CS1859 CS1883 CS1810 CS1833 CS1832 CS1856] |
| Hap_388: 2 | [CS1879 CS1818] |
| Hap_389: 5 | [CS1151 CS1155 CS1152 CS1156 CS1146] |
| Hap_390: 3 | [CS1822 CS1864 CS1802] |
| Hap_391: 5 | [CS1103 CS1826 CS1827 CS1828 CS1825] |
| Hap_392: 1 | [CS1808] |
| Hap_393: 1 | [CS1888] |
| Hap_394: 2 | [CS1124 CS1127] |
| Hap_395: 3 | [CS1140 CS1889 CS1890] |
| Hap_396: 1 | [CS1892] |
| Hap_397: 1 | [CS1797] |
| Hap_398: 1 | [CS1854] |
| Hap_399: 1 | [CS1145] |
| Hap_400: 1 | [CS1851] |
