## Supplementary Table 8a for "Initial insights into the genetic epidemiology of SARS-CoV-2 isolates from Kerala suggest local spread from limited introductions"

| Variant | Occurrence | Frequency | Total Read Count | Reference Base Count | Alternate Base Count | Percentage of variation |
|---|---|---|---|---|---|---|
| **14120C>T** | 1 | 0.56% | 13478 | C - 5 | T - 13472 | 100% |
| | | | 2398 | C - 0 | T - 2398 | 100% |
| | | | 880 | C - 0 | T - 880 | 100% |
| | | | 444 | C - 0 | T - 444 | 100% |
| | | | 11 | C - 0 | T - 11 | 100% |
| | | | 23 | C - 0 | T - 23 | 100% |
| | | | 853 | C - 0 | T - 853 | 100% |
| | | | 3646 | C - 0 | T - 3646 | 100% |
| **22444C>T** | 8 | 4.469%) | 115 | C - 0 | T - 115 | 100% |
| | | | 14272 | G - 0 | C - 14272 | 100% |
| **26367G>C** | 2 | 1.12% | 7553 | G - 2 | C - 7551 | 100% |
| | | | 5502 | C - 1 | T - 5501 | 100% |
| | | | 3218 | C - 0 | T - 3218 | 100% |
| | | | 2917 | C - 0 | T - 2917 | 100% |
| | | | 218 | C - 0 | T - 218 | 100% |
| | | | 164 | C - 0 | T - 164 | 100% |
| **28854C>T** | 6 | 3.35% | 1909 | C - 0 | T - 1909 | 100% |
| | | | 171 | G - 0 | T - 171 | 100% |
| **28899G>T** | 2 | 1.12% | 2513 | G - 8 | T - 2505 | 100% |
