## Supplementary Table 8b for "Initial insights into the genetic epidemiology of SARS-CoV-2 isolates from Kerala suggest local spread from limited introductions"

| Variant | Occurrence | Frequency | Total Read Count | Reference Base Count | Alternate Base Count | Percentage of variation |
|---|---|---|---|---|---|---|
| | | | 460 | T - 1 | C - 459 | 100% |
| | | | 201 | T - 0 | C - 201 | 100% |
| | | | 282 | T - 0 | C - 282 | 100% |
| **21486T>C** | 4 | 2.24% | 16 | T - 0 | C - 16 | 100% |
| | | | 424 | T - 0 | C - 424 | 100% |
| **23719T>C** | 2 | 1.12% | 86 | T -0 | C - 86 | 100% |
| **25281G>A** | 1 | 0.56% | 4771 | G - 2 | A - 4769 | 100% |
| | | | 139 | C - 0 | A - 139 | 100% |
| | | | 62 | C - 0 | A - 62 | 100% |
| | | | 58 | C - 1 | A - 57 | 100% |
| | | | 83 | C - 0 | A - 83 | 100% |
| **27643C>A** | 5 | 2.79% | 175 | C - 0 | A - 175 | 100% |
