## Supplementary figures and images for "Initial insights into the genetic epidemiology of SARS-CoV-2 isolates from Kerala suggest local spread from limited introductions"

### Supplementary Figure 1

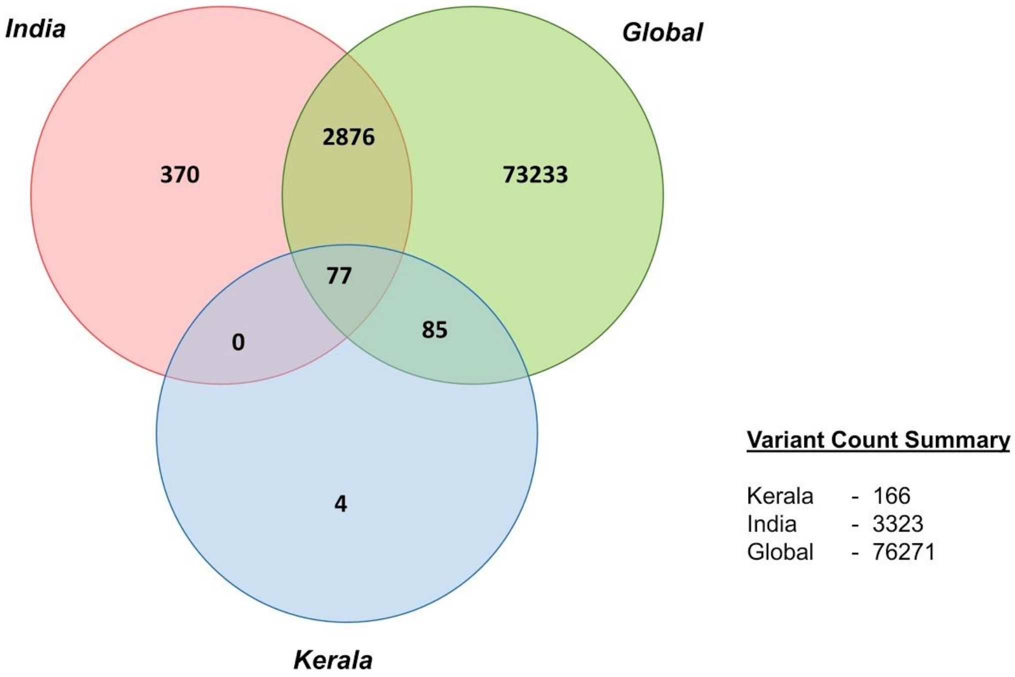


**Supplementary Figure 1.** Distribution of genetic variants across datasets
